# Lack of pulmonary fibrogenicity and carcinogenicity of titanium dioxide nanoparticles in 26-week inhalation study in rasH2 mouse model

**DOI:** 10.1101/2021.12.23.473959

**Authors:** Shotaro Yamano, Tomoki Takeda, Yuko Goto, Shigeyuki Hirai, Yusuke Furukawa, Yoshinori Kikuchi, Tatsuya Kasai, Kyohei Misumi, Masaaki Suzuki, Kenji Takanobu, Hideki Senoh, Misae Saito, Hitomi Kondo, Yumi Umeda

## Abstract

**Background:** With the rapid development of alternative methods based on the spirit of animal welfare, the publications of animal studies evaluating endpoints such as cancer have been extremely reduced. There have been no systemic inhalation exposure studies of titanium dioxide nanoparticles (TiO_2_ NPs) using CByB6F1-Tg(HRAS)2Jic (rasH2) 26-week study mice model for detecting carcinogenicity.

**Methods:** Male and female rasH2 mice were exposed to 2, 8 or 32 mg/m^3^ of TiO_2_ NPs for 6 hours/day, 5 days/week for 26 weeks using a whole-body inhalation exposure system, with reference to the Organization for Economic Co-operation and Development principles of Good Laboratory Practice. All tissues including lungs, and blood were collected and subjected to biological and histopathological analyses. Additionally, Ki67 positive index were evaluated in mice lung alveolar epithelial type 2 cell (AEC2).

**Results:** This study established a stable method for generating and exposing TiO_2_ NPs aerosol, and clarified the dose-response relationship by TiO_2_ NPs inhalation to rasH2 mice. TiO_2_ NPs exposure induced deposition of particles in lungs and mediastinal lymph nodes in a dose-dependent manner in each exposure group. Additionally, alveolar inflammation was only observed in 32 mg/m^3^ exposure group in both the sexes. Exposure to TiO_2_ NPs, as well as other organs, did not increase the incidence of lung tumors in any group, and pulmonary fibrosis and pre-neoplastic lesions were not observed in all groups. Finally, the cell proliferative activity of AEC2 was examined, and it was not increased by exposure to TiO_2_ NPs.

**Conclusions:** This is the first report showing the lack of pulmonary fibrogenicity and carcinogenicity (no evidence of carcinogenic activity) of TiO_2_ NPs in 26-week inhalation study in rasH2 mice exposed up to 32 mg/m^3^, which is considered to be a high concentration. Macrophages undergoing phagocytosis due to TiO_2_ NPs exposure formed inflammatory foci in the alveolar regions of exposed mice but did not develop fibrosis or hyperplasia or tumors. Moreover, the cell proliferative ability of AEC2 in lesions was not increased. In addition, no carcinogenicity was observed for any organs other than the lungs in this study.

## Background

The industrial production of titanium dioxide (TiO_2_) as a white pigment has been ongoing for last 100 year along with the production of TiO_2_ nano particles (NPs) with particle size in one order of magnitude smaller than that of pigment-grade for last 40 years. In Japan, TiO_2_ is approved for use in pharmaceuticals, food additives and cosmetics, and no serious health hazards have been reported till date. Pigment-grade TiO_2_ particles are widely used in paints due to their high refractive index, which gives them a bright and natural white color [1]. TiO_2_ NPs are of use when properties such as transparency and maximum ultraviolet scavenging potential are desired in sunscreens or for photocatalyst functions in the environment-purification systems [2]. There are many types of TiO_2_ depending on the combination of particle size, crystal structures, and surface modification. TiO_2_ occurs in nature in three mineral crystal structures: anatase, rutile, and brookite [3, 4]. A lot of progress has been made in development of technology for surface processing of TiO_2_ particles to give them additional characteristics [5]. Due to wider spectrum of use of TiO_2_, people are exposed to TiO_2_ through many different exposure pathways.

In recent years, the debate on the carcinogenic classification of TiO_2_ has become heated, especially in Europe. For the oral administration route, some recent studies have reported the induction of epithelial hyperplasia in the intestine of rats and mice after subacute to subchronic exposure to food-grade TiO_2_ (E171) [6–8]. However, many other oral administration studies including 2 year carcinogenicity study did not find intestinal tumor induction [9–11]. Therefore, these findings are still controversial [12]. One of the possible reasons for this is lack of long-term animal studies which could have determined toxicity of TiO_2_ exposure in terms of tumorigenesis, and carcinogenicity induced due to exposure. The available data is based on overestimated short-term animal studies and specific in vitro alternative studies. Previously conducted systemic inhalation studies in rats have reported little to significant carcinogenicity of TiO_2_ to the lungs [13] or significant carcinogenicity at high doses [14, 15]. Based on the positive results of these rat studies, the International Agency for Research on Cancer (IARC) has determinedTiO_2_ as *possibly* carcinogenic to humans, Group 2B [16]. However, there is no carcinogenicity reported for other animals including mice and hamsters [17, 18]. This emphasizes the need to assess the carcinogenicity of TiO_2_ in other animal models to further establish the carcinogenic effect of TiO_2,_ if any. Chronic inhalation studies have been decreasing worldwide in recent years due to cost, equipment, and animal welfare issues. Therefore, there is an urgent need to introduce an experimental model that can evaluate chronic toxicity, especially carcinogenicity, more rapidly and accurately.

CByB6F1-Tg(HRAS)2Jic (rasH2) transgenic mouse model has been developed as an alternative to the long-term studies (1.5years-lifetime) to predict the carcinogenic potential of chemicals [19]. This mouse model is characterized by multiple copies of the human c-Ha-ras oncogene and promoter in its genome [19]. The results of 6-month carcinogenicity studies with food additives, existing carcinogens, and oral drug candidates have demonstrated that the rasH2 mice are more sensitive to genotoxic as well as non-genotoxic carcinogens than a p53 heterozygous mouse model [20–23]. Therefore, rasH2 mouse model is clearly useful for detecting carcinogenicity in as short a time as six months, although the cost is high, more than $200,000, to study one test substance, including pilot studies. rasH2 mice have been used in 26-week carcinogenesis studies of various chemicals administered transdermally, intravenously, or orally. However, there is only one report of a whole-body inhalation exposure study in which the effects of exposure to mainstream tobacco smoke on lung tumors were examined [17]. We have already conducted systemic inhalation exposure studies of various gaseous substances using the rasH2 mouse model, and have confirmed the usefulness of this model (study reports of the Ministry of Health, Labour and Welfare, unpublished data).In the present study, we utilized this experience to conduct a 26-week systemic inhalation exposure study of TiO_2_ NPs using rasH2 mice and comprehensively evaluate the incidences of tumor development in various organs. In addition, the adverse outcome pathways (AOPs), a conceptual framework that uses existing knowledge of events at the molecular level [initiating event (IE)] and links it to an adverse outcome (AO) via key events (KEs), is useful for understanding of toxicity relationship [24]. Postulated AOP of TiO_2_ related to carcinogenicity after inhalation exposure is well summarized in previous report [18]. The particle-laden macrophages linger in the alveolar air spaces and cause impaired clearance (IE), which leads to persistent inflammation (KE3), causing alveolar epithelial type 2 cell (AEC2) proliferation (KE6) and the development of bronchiolo-alveolar hyperplasia (KE7), a preneoplastic lesion. This further leads to the induction of lung tumor (AO). In addition, although in the framework of the AOP for TiO_2_, there is no KE for lung fibrosis [18], lung fibrosis has been discussed as a KE in inhalation toxicity NPs containing TiO_2_ [25–27]. Therefore, with reference to this evidence, we also evaluated the various KEs and fibrosis.

## Results

### Stability of aerosol generation and mass concentration and particle size distribution of TiO_2_ NPs in the inhalation chamber

The mass concentration of TiO_2_ NPs aerosol in the inhalation chamber has been shown in Figure S4. Each TiO_2_ NPs concentration was nearly equal to the target concentration over the 26-week exposure period (Fig. S4). The mass median aerodynamic diameters (MMAD) and geometric standard deviation (σg) of the TiO_2_ NPs aerosol were within 0.8-1.0 μm and 2.0-2.1, respectively, and were similar for all TiO_2_ NPs-exposed groups (Fig. 4). Morphological observations by scanning electron microscope (SEM) confirmed that the TiO_2_ NPs generated in the chamber did not highly aggregated immediately (Fig. S4). These data indicate that TiO_2_ NPs aerosols were generated stably during the 26-week exposure period.

### Cytology and biochemistry of plasma, and organ weight

In all TiO_2_ NP-exposed mice, neither exposure-related mortality nor respiratory clinical signs were observed throughout the study (Fig. 1A, B). There was significant increase in the final body weight in both males and females in the 8 mg/m^3^ exposure group compared to the control group (Fig. 2A, B), but the rate of increase was very less. Males in the 32 mg/m^3^ exposure group showed a significant decrease in the red blood cell count and hematocrit value and a significant increase in MCV and MCH compared to the control group (Table S1). From these results, it was concluded that this group was mildly anemic. White blood cell counts at 32 mg/m^3^ in males and females showed a decreasing trend in males and a significant decrease in females (Table S1). Statistically significant changes were also observed in MCV and MCH, but none showed concentration correlation, and therefore were not considered to be the toxic effects (Table S1). In addition, TiO_2_ NP significantly altered several other biochemistry parameters, such as glucose and lactate dehydrogenase (LDH) (Table S2). However, most of the parameters did not show any significant correlation with the exposure concentration, suggesting low toxicological significance. TiO_2_ NP concentration-dependent increase in the lung weight was observed in both males and females (Fig. 2C-F). Although TiO_2_ NP also caused significant increase or decrease in several other organ weights, including liver, most of the changes were slight (approximately 10%) or did not correlate with the exposure concentration (Tables S3 and S4). Only in the 32 mg/m^3^group of both males and females, multiple white spots were observed in the lungs, while there were no gross findings in other organs.

**Figure 1.**
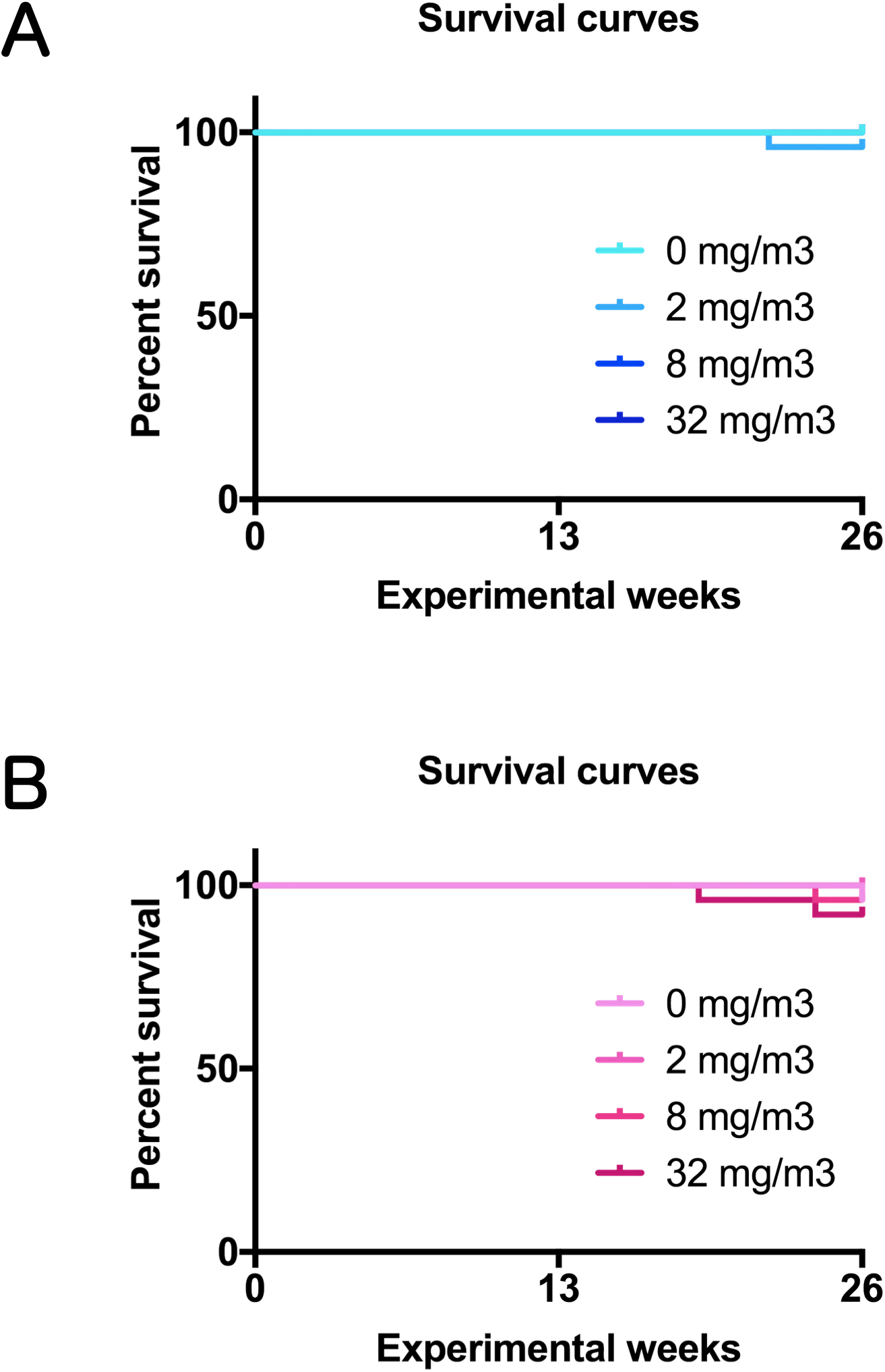
Survival curves of rasH2 mice which inhaled titanium dioxide nanoparticles (TiO_2_ NPs) (2, 8 or 32 mg/m^3^, 6 hours/day, 5 days/week, 26 weeks). A: male and B: female.

**Figure 2.**
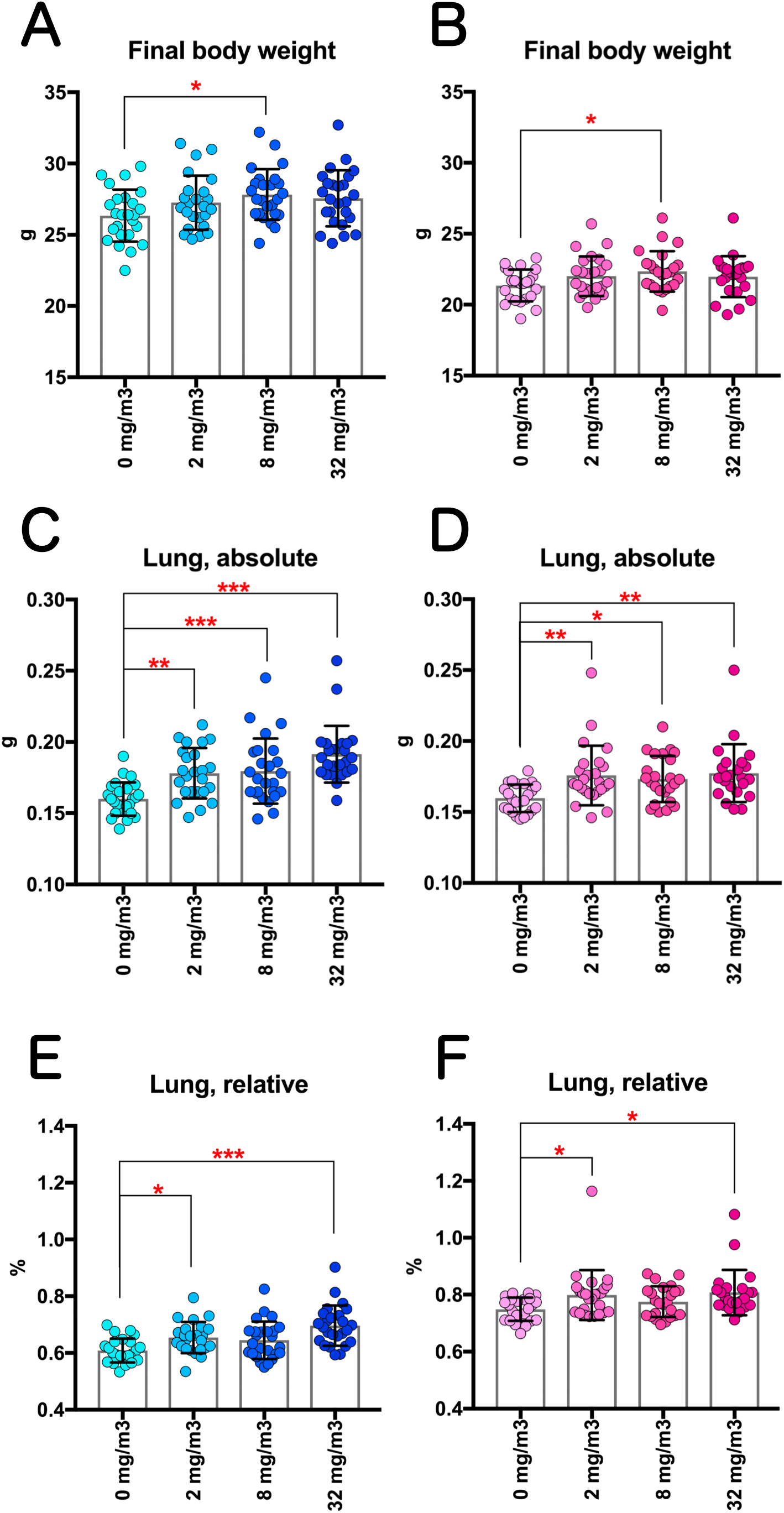
Changes in the final body weights and the lung weight of rasH2 mice following inhalation exposure to TiO_2_ NPs (2, 8 or 32 mg/m^3^, 6 hours/day, 5 days/week, 26 weeks). Final body weights in male (A) and female (B) mice, and absolute lung weights in male (C) and female (D) mice were measured at sacrifice, and the relative lung weights of males (E) and females (F) were calculated as a percentage of body weight. Dunnett’s multiple comparison test was used to compare with the age-matched control (0 mg/m^3^) groups: \**p*<0.05, \*\**p*<0.01 and \*\*\**p*<0.001.

### Histopathological findings excluding the lung and mediastinal lymph node

Except for the lungs and mediastinal lymph nodes affected by TiO_2_ inhalation, histopathological findings of nasal cavity, liver, stomach, pancreas, kidney, urinary bladder, spleen, pituitary gland, thyroid, parathyroid, adrenal, testis, epididymis, ovary, uterus, brain, harderian gland, peripheral nerve, cartilage, subcutis, pleura, mediastinum and peritoneum have been shown in Tables 1 and S5. Inhalation exposure to TiO_2_ NPs did not affect the incidence of non-neoplastic and neoplastic lesions in these organs, indicating the toxic effects of TiO_2_ NPs exposure were limited only to the alveolar region of the lungs, and were not carcinogenic to any organ in this study.

**Table 1.**
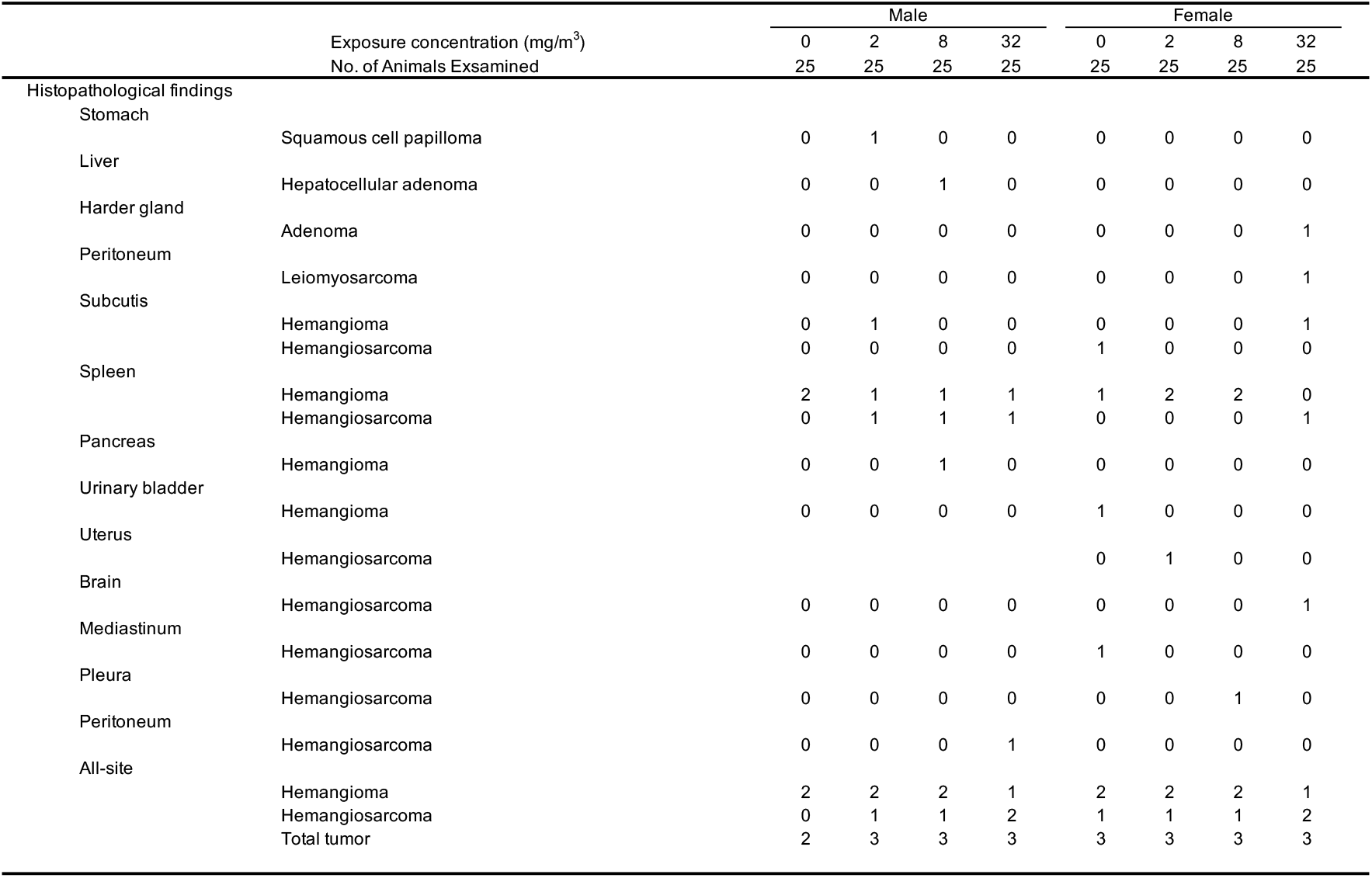
Incidence of the histopathological findings of neoplastic lesions in the all organs excluding the lung and mediastinal lymph nodes.

### Histopathological examination for lung and mediastinal lymph node

Representative microscopic photograph of the lungs has been shown in Figs. 3, S1 and S2, and histopathological findings for the lung and mediastinal lymph nodes have been summarized in Table 2. TiO_2_ NP exposure induced various particle-laden macrophage-associated pulmonary lesions. Deposition of the particles, commonly observed after the inhalation exposure to the particles, was observed in all the mice exposed to TiO_2_ NPs (Fig. 3B). The lesion intensity was found to be concentration-dependent (Table 2). The extrapulmonary ejection of TiO_2_ NPs through the muco-ciliary escalator was observed in the all exposed groups, including the 32 mg/m^3^ group (Fig. 3C). In addition, inflammatory cell infiltration with particle-laden macrophage in the alveolar region was caused only by 32 mg/m^3^ exposure in both of the sexes (Fig. 3D and Table 2), and particle-laden macrophages in the lesions phagocytosed TiO_2_ NPs to the extent that the nuclei were invisible. The lesions were difficult to be observed as a focus in the formalin-inflated left lung (Fig. S2A-B), indicating that the lesions move easily in the alveoli by inflation. On the other hand, the BALT swelling due to exposure to TiO_2_ NPs was not observed in any mice (Figs. 3, S1and S2).

**Table 2.** Incidence and integrity of the histopathological findings of the lung and mediastinal lymph nodes.

|  |  | Male |  |  |  | Female |  |  |  |
| --- | --- | --- | --- | --- | --- | --- | --- | --- | --- |
| Exposure concentration (mg/m<sup>3</sup>) |  | 0 | 2 | 8 | 32 | 0 | 2 | 8 | 32 |
| No. of Animals Examined |  | 25 | 25 | 25 | 25 | 25 | 25 | 25 | 25 |
| Histopathological findings |  |  |  |  |  |  |  |  |  |
| Mediastinal lymph node |  |  |  |  |  |  |  |  |  |
|  | Deposit of particle | 0 | 0 | 0 | 13***<br><2> | 0 | 0 | 0 | 8**<br><2> |
| Lung |  |  |  |  |  |  |  |  |  |
| Non-neoplastic |  |  |  |  |  |  |  |  |  |
|  | Deposit of particle | 0 | 25***<br><1> | 25***<br><2> | 25***<br><3> | 0 | 25***<br><1> | 25***<br><2> | 25***<br><3> |
|  | Inflammatory cell infiltration with particle-laden macrophage | 0 | 0 | 0 | 13***<br><1> | 0 | 0 | 0 | 8**<br><1> |
|  | Fibrosis, alveolar interstitium | 0 | 0 | 0 | 0 | 0 | 0 | 0 | 0 |
|  | Hyperplasia, Bronchiolo-alveolar | 0 | 0 | 0 | 0 | 0 | 0 | 0 | 0 |
| Neoplastic |  |  |  |  |  |  |  |  |  |
|  | Adenoma, Bronchiolo-alveolar | 1 | 4 | 1 | 4 | 0 | 0 | 0 | 2<sup>#</sup> |
|  | Carcinoma, Bronchiolo-alveolar | 0 | 0 | 0 | 1 | 1 | 1 | 0 | 0 |
|  | Total tumor | 1 | 4 | 1 | 5 | 1 | 1 | 0 | 2 |

Note: Values indicate number of animals bearing lesions. The value in angle bracket indicate the average of severity grade index of the lesion. The average of severity grade is calculated with a following equation. $\Sigma(\text{grade} \times \text{number of animals with grade}) / \text{number of affected animals}$ . Grade: 1, slight; 2, moderate; 3, marked; 4, severe. Significant difference: \*, p<0.05; \*\*, p<0.01; \*\*\*, p<0.001 by Chi square test compared with each cont. #: p<0.05 by Peto and Cochran-Armitage trend test.
Note: Values indicate number of animals bearing lesions. The value in angle bracket indicate the average of severity grade index of the lesion. The average of severity grade is calculated with a following equation. $\Sigma(\text{grade} \times \text{number of animals with grade}) / \text{number of affected animals}$ . Grade: 1, slight; 2, moderate; 3, marked; 4, severe. Significant difference: \*, p<0.05; \*\*, p<0.01; \*\*\*, p<0.001 by Chi square test compared with each cont.

**Figure 3.**
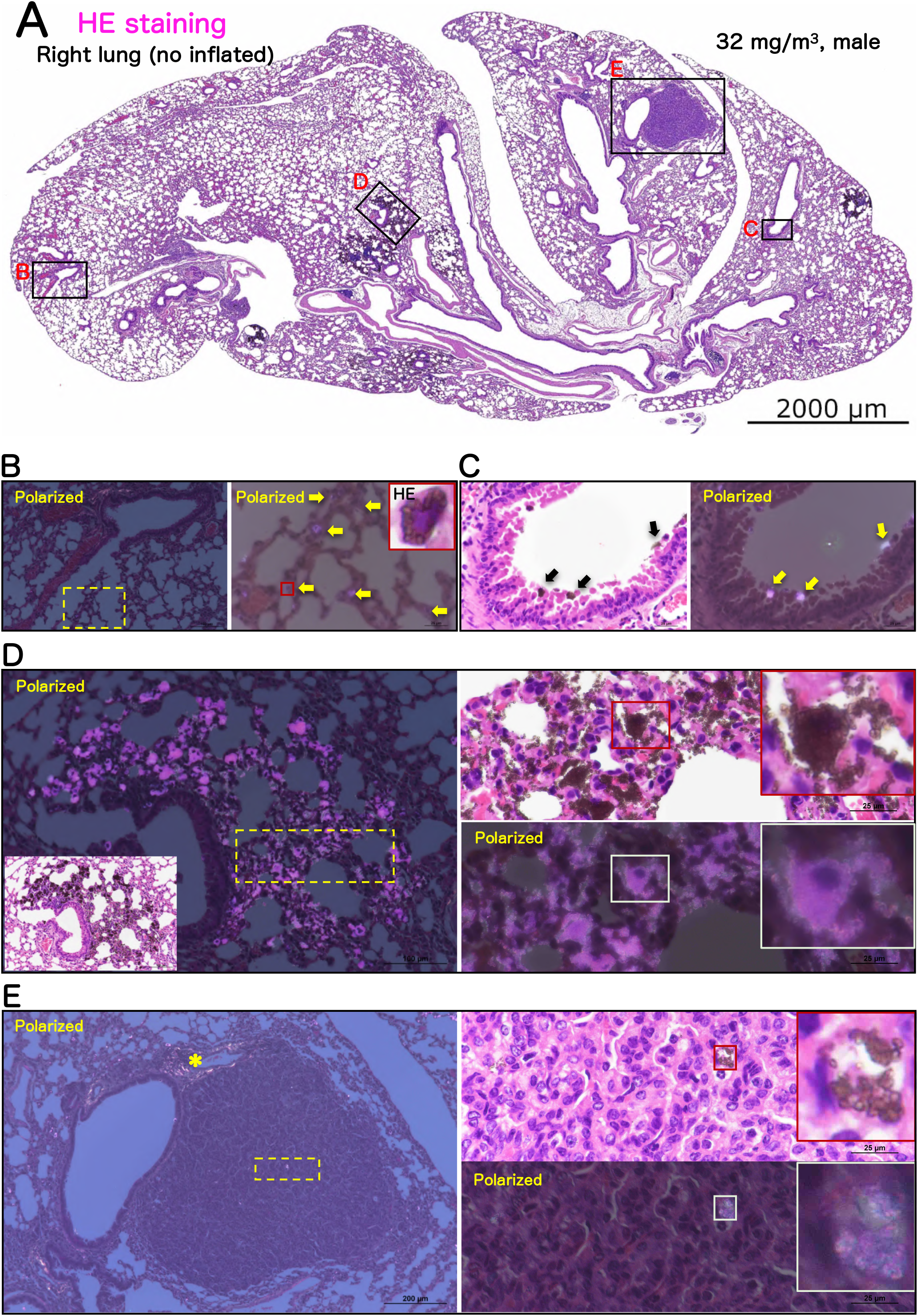
Representative microscopic photographs of the male rasH2 mouse right lungs after inhalation exposure to TiO_2_ NPs (32 mg/m^3^). The right lung was not injected with formalin through the bronchus into the lung, and formalin immersion fixation was performed after the lung was removed. The lungs were stained with hematoxylin and eosin (HE) and all images were taken with HE or polarized light microscope. TiO_2_ NPs, naked or phagocytosed by macrophages, were observed mainly in the alveolar region and were black in HE staining or pink in polarized light microscope. A typical loupe image (A) of the entire right lungs and magnified images of each lesions (B-E) have been shown. Deposit of particle has been observed in single particle-laden macrophage or naked particle (B), particles in the process of being eliminated by the mucociliary escalator were observed on the bronchial mucosa (C). Inflammatory cell infiltration with particle-laden macrophage (Inflammatory foci) was observed as a focus under low magnification (left side of panel D), and a large amount of particles were observed to have clumped together under polarized light. Macrophages present in the lesion phagocytosized the particles until the nucleus was not visible (right side of panel D). Representative image of bronchiolo-alveolar adenoma (E). Histopathologically, the morphology of the tumors observed in the TiO_2_ NPs -exposed group was not different from that observed in the control group, and the amount of particles or particle-laden macrophages observed in the tumors was much lower than that in the inflammatory foci (right side of panel E). *: collagen fibers of axal interstitium.

To examine the changes in the fiber volume, Masson’s trichrome staining in the lungs was performed (Figs. 4 and S3). The results showed that, in the lungs of the 32 mg/m^3^ exposure group, there was no increase in Masson stain-positive areas in the alveolar septa and perivascular interstitium within the lesions (Fig. 4B) when compared to the lesion surrounding area (Fig. 4C) of exposed lung and peribronchiolar and subpleural alveolar regions of control lung (Fig. S3). These results indicated that TiO_2_ NPs do not cause by fibrosis of the alveolar interstitium in this study model (Table 2).

**Figure 4.**
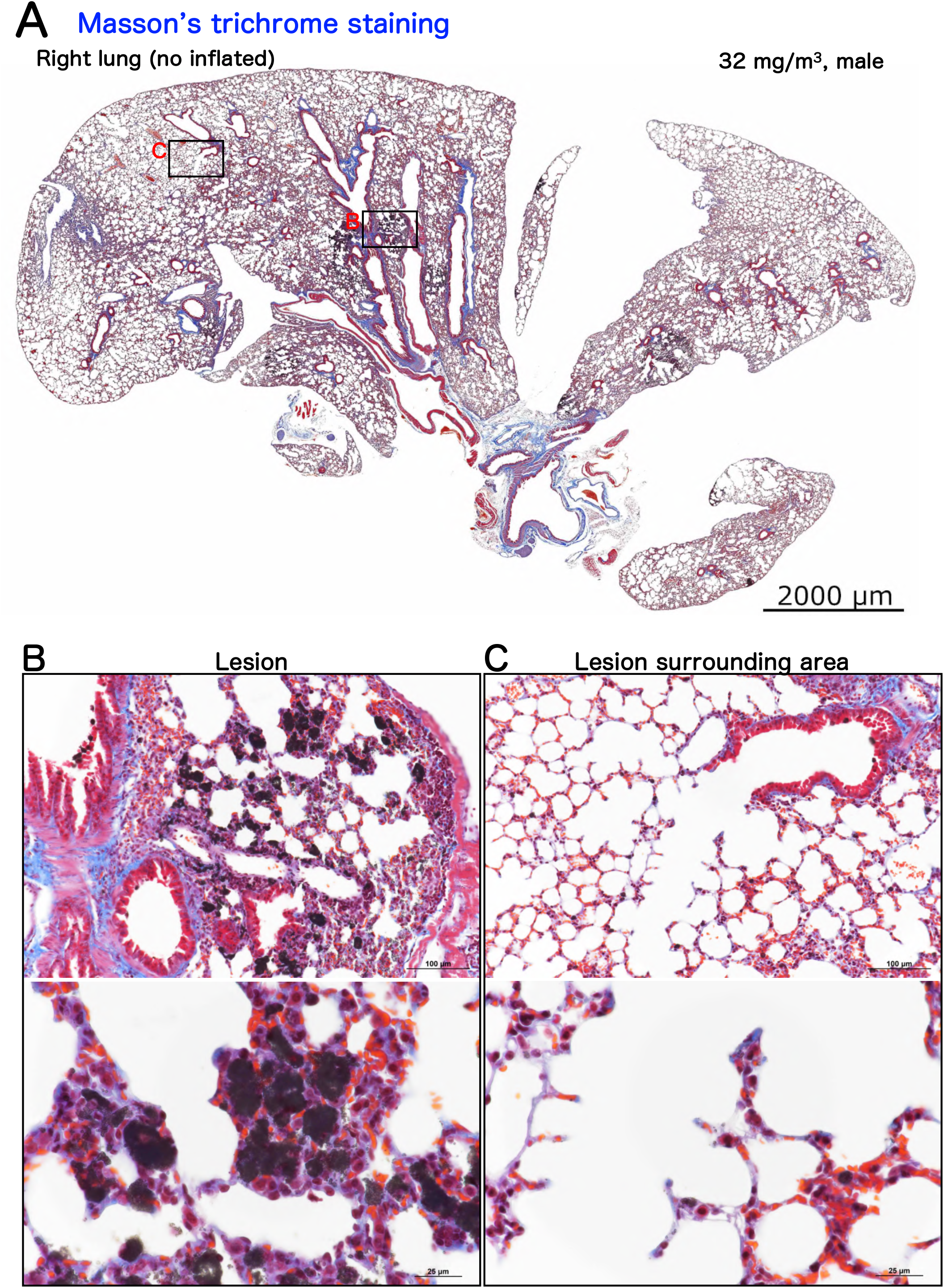
Representative microscopic photographs of Masson’s trichrome staining of male rasH2 mouse right lungs after inhalation exposure to TiO_2_ NPs (32 mg/m^3^). A typical loupe image (A) of the entire right lungs and magnified images of lesion (B) and lesion surrounding area (C) have been shown. Regardless of the presence or absence of a lesion, no expansion of the blue area was observed in the alveolar septa.

We further examined the occurrence of lung tumors, bronchiolo-alveolar adenomas (Fig. 3E) and carcinomas, in both the sexes. There was no continuity/co-localization with inflammatory lesions in any of the tumors exposed to TiO_2_ NPs, and the number of infiltrating particle-laden macrophages in the tumors was limited (Fig. 3E). Histopathological growth pattern and characteristics of the tumor stroma were not different from that of lung tumors in the control group. There was no significant increase in the incidence of bronchiolo-alveolar adenoma, carcinoma, or total tumor in the males exposed groups compared to the control group (Table 2). On the other hand, in females, a significant increase was observed in the incidence of bronchiolo-alveolar adenoma by Peto and Cochran-Armitage trend tests (Table 2). However, no significant difference was observed in the incidence of bronchiolo-alveolar carcinoma and total tumor in any of the TiO_2_ exposure group. Furthermore, TiO_2_NPs exposure did not cause hyperplasia, a preneoplastic lesion of lung tumor (Table 2). Taken together, these results indicated that no fibrogenicity and carcinogenicity was caused to the lungs of rasH2 mice exposed to TiO_2_ NPs of both the sexes. In mediastinal lymph nodes, deposition of particles was observed in both the sexes of the 32 mg/m^3^ exposure group, with a significant increase (Table 2). However, this finding was not considered to be a toxic effect because there were no grossly enlarged lymph nodes or tissue findings such as lymphoid hyperplasia.

### Cell proliferation ability in Alveolar epithelial type 2 cells in mice lung

Similar to the previously reported AOP of TiO_2_ [18], inhalation exposure to TiO_2_ induced one of KEs, persistent inflammation (KE3), although neither bronchiolo-alveolar hyperplasia (KE7) nor fibrosis (AO-Extra) or tumors (AO) associated with inflammatory foci (KE3) were observed in this study. Next, to investigate another KE, the proliferation ability of AEC2 (KE6), we performed double staining for Ki67, a cell proliferation marker, and Lysophosphatidylcholine acyltransferase 1 (LPCAT1), an AEC2 marker.

Slightly increasing trend was observed in the LPCAT1-positive AEC2 in the lesions of both the sexes in the 32 mg/m^3^ group compared to both the alveolar area at 0 mg/m^3^ and the lesion surrounding area in the 32 mg/m^3^ group (Fig. 5A). However, the Ki67 positive index showed no significant increase in the percentage of Ki67 positive AEC2 in the lesion in both the sexes (Fig. 5B). These results indicated that the cell proliferative ability of AEC2 in lesions was not increased by TiO_2_ NPs inhalation under the conditions of this study.

**Figure 5.**
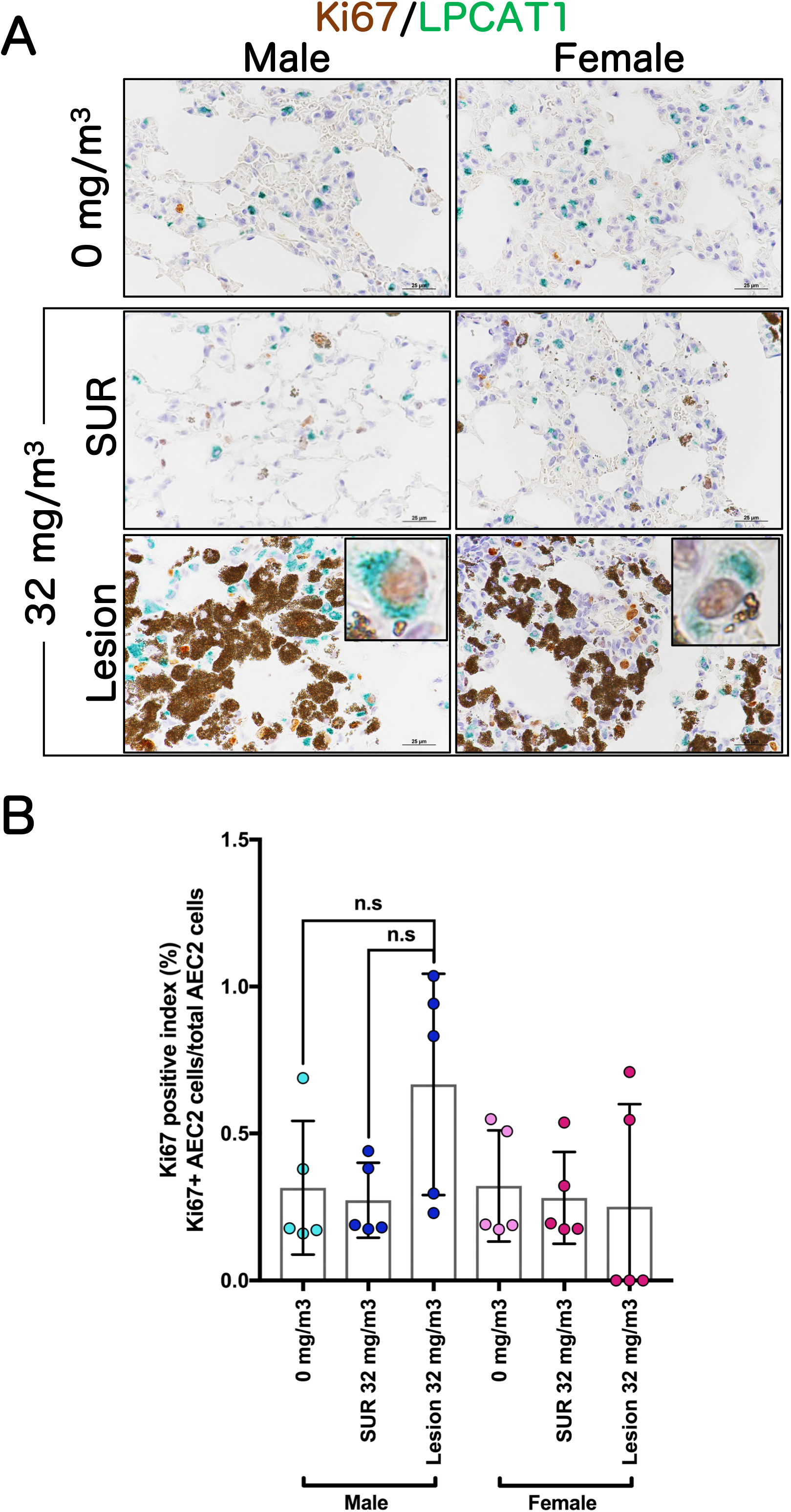
Examination for cell proliferative activity in alveolar epithelial type 2 cell (AEC2). Representative immunohistochemical staining images of Ki67, a cell proliferation marker, and Lysophosphatidylcholine acyltransferase 1 (LPCAT1), an AEC2 marker (A). The Ki67-positive index in AEC2 was calculated as the percentage of Ki67 and LPCAT1 double positive cells to total LPCAT1-positive cells. Abbreviations: Lesion: inflammatory foci, SUR: lesion-surrounding tissue and ns: not significant.

**Figure 6.**
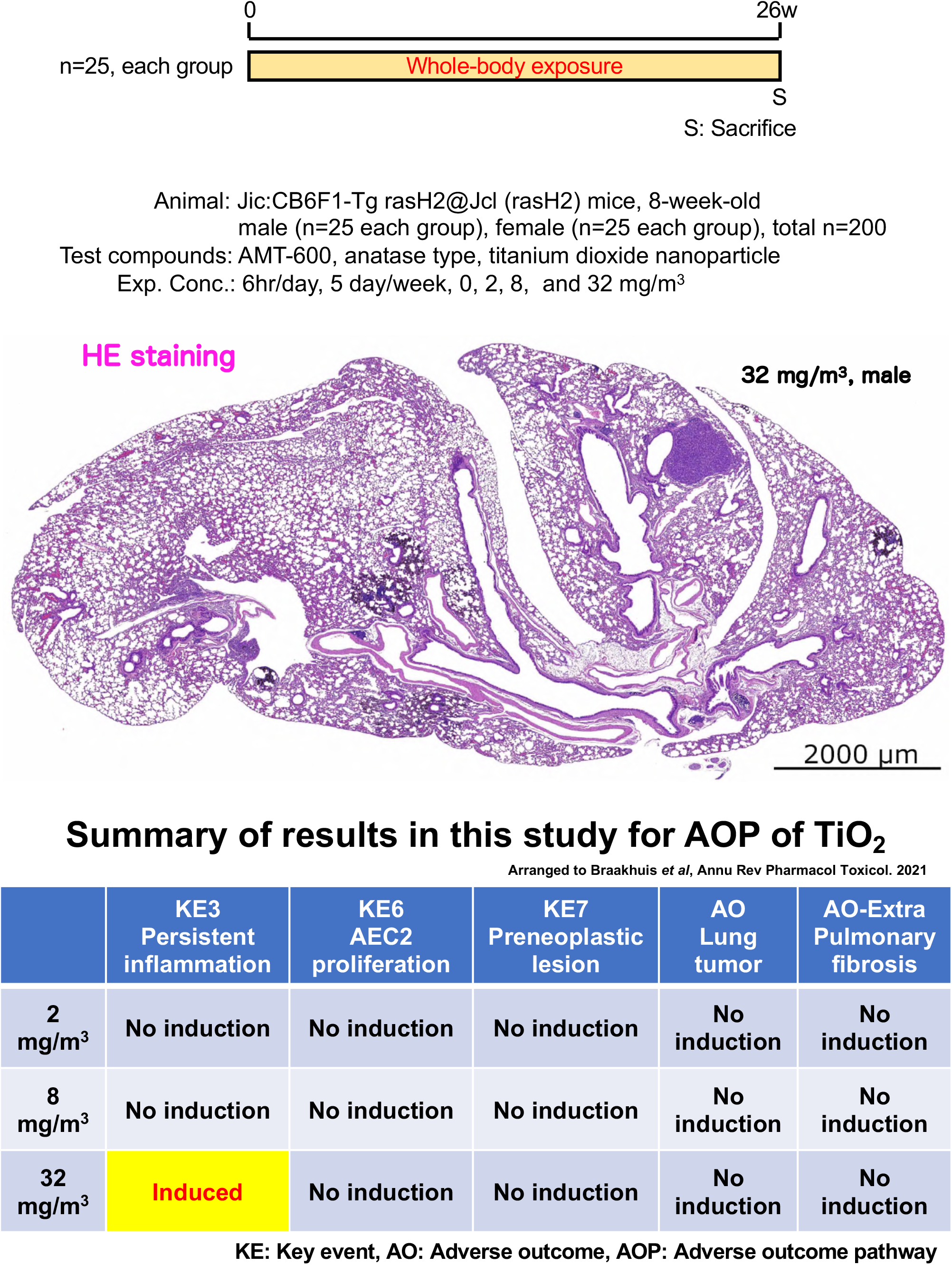
Graphical abstract of this study.

## Discussion

Herein, we examined the carcinogenicity of TiO_2_ NPs in 26-week inhalation study using rasH2 mouse model. The results showed that no significant carcinogenicity was observed in all the organs including the lungs in both sexes at exposures up to 32 mg/m^3^, which is considered to be a high concentration. As for toxic effects in the lungs is concerned, phagocytosis was observed in macrophages after exposure to TiO_2_ NPs and neutrophils formed inflammatory foci in the alveolar region, but these did not progress to hyperplasia and tumors but also fibrosis. In February 2006, IARC Monographs Working Group re-evaluated the carcinogenic hazards of TiO_2_ to humans and this was published in the IARC Monographs [16]. The results from the rodent carcinogenic studies with TiO_2_ were considered to provide *sufficient* evidence for its carcinogenicity by the Working Group and TiO_2_ was established as *possibly* carcinogenic to humans, Group 2B [16]. This was based on the findings of the long-term systemic inhalation exposure studies using pigment-grade TiO_2_ [14] and TiO_2_ NPs [15] and serial intratracheal administration studies [28] that showed lung carcinogenicity in rats. However, all the data showing carcinogenicity of TiO_2_ are limited to rats. No lung carcinogenicity has been reported in other experimental animal models such as mice and hamsters [22, 24]. Moreover, long-term TiO_2_ administration studies via various routes, including oral, intraperitoneal, and subcutaneous in any experimental animal have not shown carcinogenicity of TiO_2_ in any organ [9,30,31]. Therefore, the data obtained in the present study using rasH2 mice is consistent with these previous reports.

In the present study, KEs that lead to the induction of tumors were examined, as no lung tumors (AO) were found to be induced. Multifocal inflammatory cell infiltrating foci with particle-laden macrophages was observed in the 32 mg/m^3^ exposure groups of both the sexes. Based on this finding, it was postulated that KE3 (persistent inflammation) was occurring in these groups. Nevertheless, not only the development of pre-neoplastic lesions (KE7), but also the increase of proliferative activity in inflammatory foci in AEC2 (KE6), which is known to be the originating cell of preneoplastic lesions, was not observed in any of the exposure groups. Previous reports on inhalation exposure studies in mice have shown that inflammation is induced after inhalation exposure to high concentrations of TiO_2_ (>10 mg/m^3^) [32–36]. In coherence with these findings, KE3 was induced only in the 32 mg/m^3^ group of both the sexes in this study, but not at concentrations below 8 mg/m^3^.Yu *et al* [33] conducted a 4-week systemic inhalation exposure study with exposure concentrations up to 10 mg/m^3^ in A/J Jms Slc mice, a mouse strain with a predilection for lung tumors. They reported that not only inflammation (KE3) but also hyperplasia (KE7) was induced in the lungs as observed histopathologically. They also found that proliferating cell nuclear antigen (PCNA) (KE6) increased in a dose-dependent manner in WB analysis of lung tissue [33]. However, the incidence and multiplicity of hyperplasia were not evaluated, and the cell types of PCNA-positive cells were also not identified. Therefore, it is controversial whether the induction of KE6 and KE7 occurred in their study. On the contrary, we examined the proliferative activity of cells limited to AEC2, which is considered to be the origin of lung tumors [37, 38], and is a more accurate strategy for evaluation of KE6 than that used by Yu *et al*. TiO_2_ is poorly soluble under normal physiological conditions and has a half-time of greater than 100 days upon short-term exposures [39–41]. Furthermore, it is known to be a substance with lower acute toxicity than copper oxide, which is highly soluble. Therefore, based on these characteristics and reports related to toxicity, TiO_2_ is classified as a poorly soluble, low toxicity (PSLT) particle [39–41]. The definition of PSLTs is still under debate, and meetings are being rigorously held worldwide on this topic [42]. For the mechanism of lung carcinogenesis of PSLT particles, the overload theory via particle-laden macrophages that continuously remain in the alveolar air spaces and the contribution of the indirect carcinogenic mechanism due to the persistent inflammatory state are mentioned [43–46]. In the present study, since both the particle-laden macrophages in the alveolar air spaces and the persistent inflammatory state in the lungs were caused by 32 mg/m^3^ exposure, it is reasonable to consider that this is a sufficient exposure concentration to assess toxicity, including carcinogenicity. However, no pulmonary tumors were observed, despite the use of rasH2 mice, a carcinogenically sensitive model. Noticeably, it has recently been reported that oral administration of E171 induces preneoplastic lesions of gastrointestinal tumors in rodents[6–8], but in this study of whole-body inhalation exposure, no gastrointestinal preneoplastic lesions, tumors and inflammatory lesions were observed. Moreover, there was no increase of collagen deposition in alveolar septa, and pulmonary fibrosis was also not observed in all the groups. Taken together, results suggested that this type of TiO_2_ NPs have little carcinogenic or fibrotic effects, at least for mice.

The present study had a limitation. Although this experiment was conducted with reference to the OECD Guidelines for the Testing of Chemicals (TG413) [47], the mouse lungs were very small compared to the rat lungs, and because priority was given to the measurement of lung weight and detailed histopathological analysis of the lungs, measurements of cytological and biochemical analyses using bronchoalveolar lavage fluid and lung burden could not be performed. This made it difficult to make quantitative comparisons of toxicity with other studies, and the contribution of overload in the toxicity mechanism could not be clearly addressed. In addition, in order to accurately evaluate the carcinogenicity of the TiO_2_ NPs used in this study to rodents and ascertain the findings, it is very important to also conduct carcinogenicity studies in rats by whole-body inhalation exposure. We are currently conducting a 13-week and 104-week systemic inhalation study in F344 rats, and the results are being compiled.

## Conclusions

This study provided first evidence, to the best of our knowledge, for the lack of pulmonary fibrogenicity and carcinogenicity (no evidence of carcinogenic activity) after exposure to TiO_2_ NPs in rasH2 mice model. We performed 26-week inhalation study in rasH2 mice exposed up to 32 mg/m^3^, which is considered to be a high concentration.

Macrophages rich in TiO2 NPs phagocytosed in the alveolar regions of exposed mice forming inflammatory foci, but did not develop into fibrosis or hyperplasia or tumors. Moreover, the cell proliferative ability of AEC2 in lesions was not increased. In addition, no carcinogenicity was observed for any organs other than the lungs in this study.

## Methods

### Materials

Anatase type TiO_2_ NP was purchased from Tayca co. (primary particle size: 30 nm). Other reagents used in the study were of the highest grade available commercially.

### Animals

6 weeks old male and female rasH2 mice were purchased from CLEA Japan, Inc. (Tokyo, Japan). Mice were housed in an air-conditioned room under a 12 hours light/12 hours dark (8:00-20:00, light cycle) photoperiod, and fed a general diet (CRF-1, Oriental Yeast Co. Ltd., Tokyo, Japan) and tap water *ad libitum*. After approximately 1-2 weeks of quarantine and acclimation, they were exposed to TiO_2_ NPs from 8 weeks of age by the procedure mentioned below. All the animal experiments were approved by the Animal Experiment Committee of the Japan Bioassay Research Center.

### Generation of TiO_2_ NP aerosol

Generation of TiO_2_ NP aerosol into the inhalation chamber was performed using our established method (cyclone sieve method) [48, 49] with some modifications. Briefly, TiO_2_ NP was fed into a dust feeder (DF-3, Shibata Scientific Technology, Ltd., Soka, Japan) to generate TiO_2_ NP aerosol, and then introduced into a particle generator (custom-made by Seishin Enterprise Co., Ltd., Saitama, Japan) ) to separate the aerosol and further supply it to the inhalation chamber. The concentration of the TiO_2_ NP aerosol in the chamber was measured and monitored by an optical particle controller (OPC; OPC-AP-600, Shibata Scientific Technology), and the operation of the dust feeder was adjusted by feedback control based on the upper and lower limit signals to maintain a steady state.

The mass concentration of TiO_2_ NP aerosol in the chamber was measured every two weeks during the exposure period. Aerosols collected on a fluoropolymer binder glass fiber filter (T60A20, φ55 mm, Tokyo Dylec, Corp., Tokyo, Japan) were weighed for each target concentration at 1, 3, and 5 hours after the start of the exposure. Using mass per particle (K-value) calculated using the measured mass results (mg/m^3^) and the particle concentration data (particles/m^3^) obtained from the OPC the particle concentration for each group during the exposure period was converted to a mass concentration. The particle size distribution and morphology of the TiO_2_ NPs were measured at first, 13th and 25th weeks of the exposure. The particle size distribution was measured using a micro-orifice uniform deposit cascade impactor (MOUDI-II, MSP Corp., Shoreview, MN). The MMAD and σg were calculated by cumulative frequency distribution graphs with logarithmic probability (Fig. S1). The TiO_2_ NP in the inhalation chamber was collected on a 0.2 μm polycarbonate filter (φ47 mm, Whatman plc, Little Chalfont, UK), and observed using SEM (SU8000, Hitachi High-Tech, Tokyo, Japan).

### 26-week inhalation study

This experiment was conducted with reference to the OECD Guidelines for Testing of Chemicals (TG413) [47]. For dosage setting, we conducted a preliminary study of 4-week inhalation exposure of TiO_2_ NPs (6.3, 12.5 25 or 50 mg/m^3^) in accordance with the OECD TG 412. A decrease in food intake was observed among the females in the 50 mg/m^3^ exposure group. Considering the increase in the lung burden due to the exposure period (from 4 to 26 weeks), target concentrations for TiO_2_ NP aerosols were at 2, 8 and 32 mg/m^3^. The exposure schedule was 6 hours per day; 5 days per week, for weeks (Fig. S7A). Two hundred mice with 25 males and 25 females in each group were transferred to the individual stainless steel cages and exposed to TiO_2_ NP for 6 hours without feeding and providing water. After daily exposure, the mice were returned to the stainless steel bedding cages and kept in groups with free access to food and water. During the study period, body weight and food consumption of mice were measured a week. At 1-4 days after the last exposure, the blood of the mice was collected under isoflurane anesthesia and euthanized by exsanguination. For histopathological analysis, all the tissues were collected from all mice in each group and fixed in 10% neutral phosphate buffered formalin solution.

### Hematological and blood chemistry tests

For hematological examination, blood samples collected at the time of each autopsy analyzed with an automated hematology analyzer (ADVIA120, Siemens Healthcare Diagnostics Inc. Tarrytown, NY). For biochemical tests, the blood was centrifuged at 3,000 rpm (2,110 × *g*) for 20 minutes, and the supernatant was analyzed with an automated analyzer (Hitachi 7080, Hitachi, Ltd., Tokyo, Japan).

### Histopathological analysis

Serial tissue sections were cut from paraffin-embedded lung specimens, and the first section (2-μm thick) was stained with H&E for histological examination and the remaining sections were used for immunohistochemical analysis. The histopathological finding terms used in this study for lesion were determined by the certified pathologists from the Japanese Society of Toxicologic Pathology, based on the finding terms adopted by International Harmonization of Nomenclature and Diagnostic Criteria for Lesions in Rats and Mice (INHAND)[50]. Pathological diagnosis was performed blindly by three pathologists and summarized after a cumulative discussion.

### Masson’s Trichrome staining

Details of the method have been described previously [51]. Briefly, the slides were deparaffinized, washed with water, and then treated with an equal volume mixture of 10% potassium dichromate and 10% trichloroacetic acid for 60 minutes at room temperature. The specimens were then washed with water and stained with Weigelt’s iron hematoxylin solution (C.I.75290, Merck-Millipore) for 10 minutes at room temperature. These slides were further stained successively with 0.8% orange G solution (C.I.16230, Merck-Millipore) for 10 minutes at room temperature, Ponceau (C.I.14700, FUJIFILM-Wako Pure Chemical Corp., Osaka, Japan), acid fuchsin (C.I.42685, Merck-Millipore), and azofloxine (C.I.18050, Chroma Germany GmbH, Augsburg, Germany) mixture for 40 minutes at room temperature, 2.5% phosphotungstic acid for 10 minutes at room temperature, and blue aniline solution (C.I.42755, Chroma Germany GmbH) under a microscope. Between staining with each solution, light washing was done with 1% acetic acid water. Thereafter, dehydration, permeabilization, and sealing were performed.

### Immunohistological multiple staining analyses

This procedure has been described in detail in a previously published study [51]. The rabbit polyclonal Ki67 antibody (ab15580) was purchased from Abcam plc (Cambridge, UK) and the mouse monoclonal LPCAT1 antibody (66044-1-lg) was purchased from proteintech (Manchester, UK). Briefly, lung tissue sections were deparaffinized with xylene, and hydrated through a graded ethanol’s series. For immunohistochemical staining, tissue sections were incubated with 0.3% hydrogen peroxide for 10 min to block the endogenous peroxidase activity. Sections were then incubated with 10% normal serum at room temperature (RT) for 10 min to block the background staining, and then incubated for 2 hr at RT with each of the primary antibodies (Ki67). After washing with PBS, the sections were incubated with histofine simple stain kit mouse MAX-PO(R) (414341, Nichirei, Tokyo, Japan) for 30 min at RT. After further washing with PBS, sections were incubated with DAB EqV Peroxidase Substrate Kit, ImmPACT (SK-4103, Vector laboratories) as brown chromogen for 2-5 min at RT for colorization. As a crucial step, after washing with dH_2_O after color detection, the sections were treated with citrate buffer at 98°C for 30 min before incubating with the next primary antibodies (LPCAT1) to denature the antibodies bound on the sections used. After that, sections were incubated with the second antibody similarly (mouse stain kit, 414321, Nichirei, Tokyo, Japan). Histogreen chromogen (AYS-E109, Cosmo Bio, Tokyo, Japan) was used for the second coloration, followed by hematoxylin staining for 30-45 seconds as a contrast stain, dehydrated and sealed. For the ki67 positive index, we counted more than 500 AEC2 in normal lungs and lesion-surrounding tissues of the 32 mg/m3 exposure group, and a total of 76 to 435 AEC2 per mouse in the lesions. The section was observed under an optical microscope ECLIPSE Ni (Nikon Corp., Tokyo, Japan) or BZ-X810 (Keyence, Osaka, Japan).

### Statistical analysis

Except in the case of incidence and integrity of histopathological lesions, the data comparisons among multiple groups were performed by one-way analysis of variance with a post-hoc test (Dunnett’s or Tukey’s multiple comparison test), using GraphPad Prism 5 (GraphPad Software, San Diego, CA). The incidences and integrity of lesions were analyzed by chi-square test using GraphPad Prism 5 (GraphPad Software, San Diego, CA). All statistical significance was set at *p* < 0.05.

## Availability of data and materials

The datasets used and analyzed during the current study are available with the corresponding authors on reasonable request.

## Ethics declarations

All animals were treated as per animal ethical protocols and all procedures were performed in compliance with the Animal Experiment Committee of the Japan Bioassay Research Center.

## Abbreviations

AEC2: Alveolar epithelial type 2 cell
AO: adverse outcome
AOP: adverse outcome pathway
σg: geometric standard deviation
HE: Hematoxylin and eosin
IARC: International Agency for Research on Cancer
IE: initiating event
KE: key event
LDH: Lactate dehydrogenase
LPCAT1: Lysophosphatidylcholine acyltransferase 1
MMAD: Mass median aerodynamic diameter
NBF: Neutral buffered formalin
PSLT: poorly soluble, low toxicity
rasH2: Jic:CB6F1-Tg ras H2@Jcl
SEM: Scanning electron microscope
SUR: lesion-surrounding tissue
TiO_2_ NPs: Titanium dioxide nanoparticles

## Acknowledgments

We would like to express our sincere gratitude to the members of Matsuzawa Kosan and Total Service for their strong support in the breeding and dissection of the animals. Finally, we would like to express our heartfelt gratitude to all the Japan Bioassay Research Center staff.

## Author information

Shotaro Yamano and Tomoki Takeda contributed equally to this work.

## Japan Bioassay Research Center, Japan Organization of Occupational Health and Safety, Kanagawa, Japan

Shotaro Yamano, Tomoki Takeda, Yuko Goto, Shigeyuki Hirai, Yusuke Furukawa, Yoshinori Kikuchi, Tatsuya Kasai, Kyohei Misumi, Masaaki Suzuki, Kenji Takanobu, Hideki Senoh, Misae Saito, Hitomi Kondo, Yumi Umeda

## Contributions

S.Y. and T.T. performed the experiments and analyzed the data. Y.G., S.H., Y.F., Y.K., K.T., M.S., S.M., S.M. and K.H. assisted with animal experiment including exposure animal care and sacrifices. K.T., H.S., Y.U., and S.Y. performed histopathological diagnoses. S.Y. and T.T. conceived, designed, and directed the study and interpreted the data. S.Y., T.T., and Y.U. drafted and revised the manuscript. All authors approved the manuscript as submitted.

## Corresponding authors

Correspondence to Shotaro Yamano or Tomoki Takeda

---

**Figure S1.**
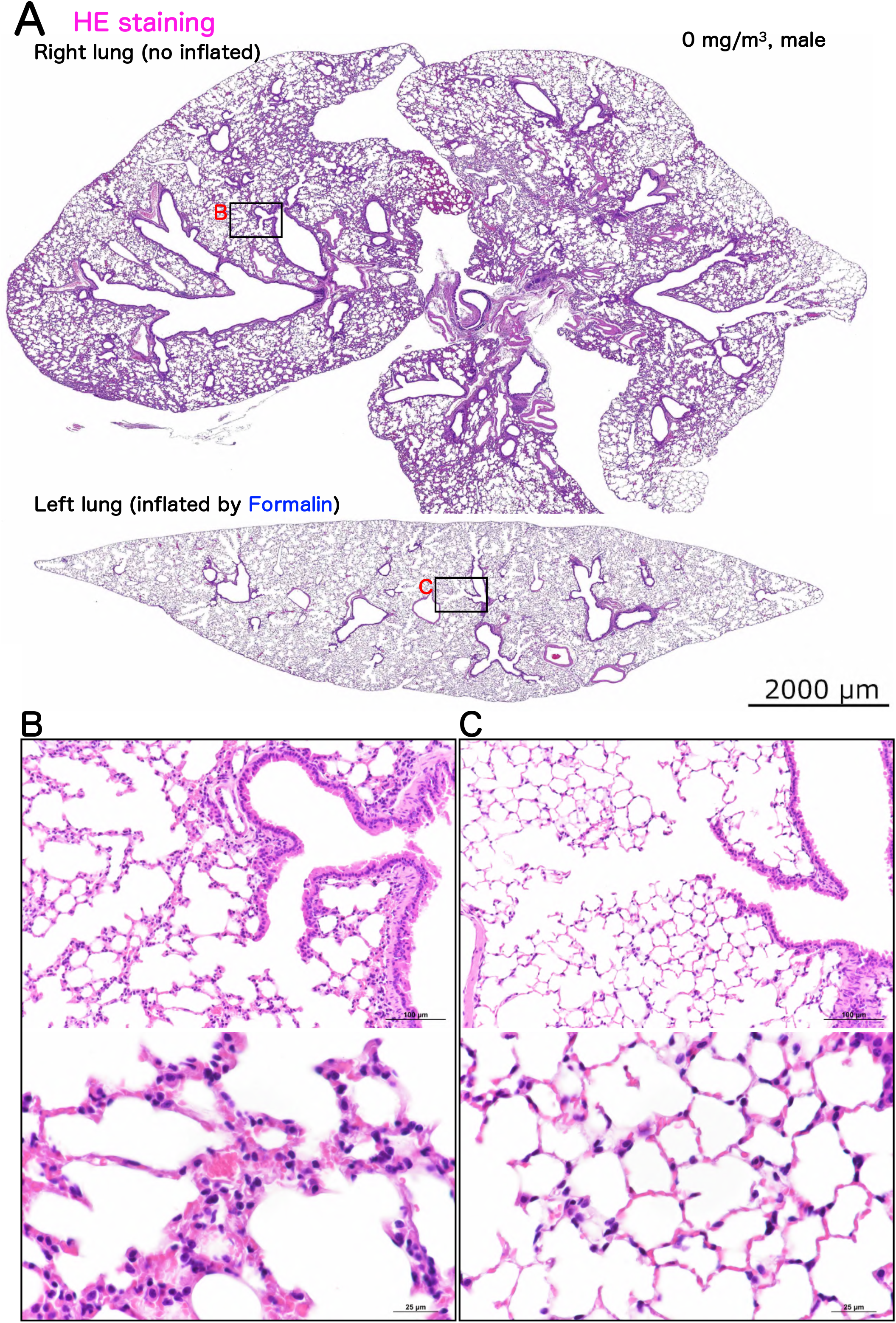
Representative microscopic photographs of lungs of a male control rasH2 mouse. The lungs were stained with hematoxylin and eosin (HE) (see the Fig. 3 legend for details). A typical loupe image (A) of the entire lungs and magnified images of normal alveolar regions (B and C) have been shown.

**Figure S2.**
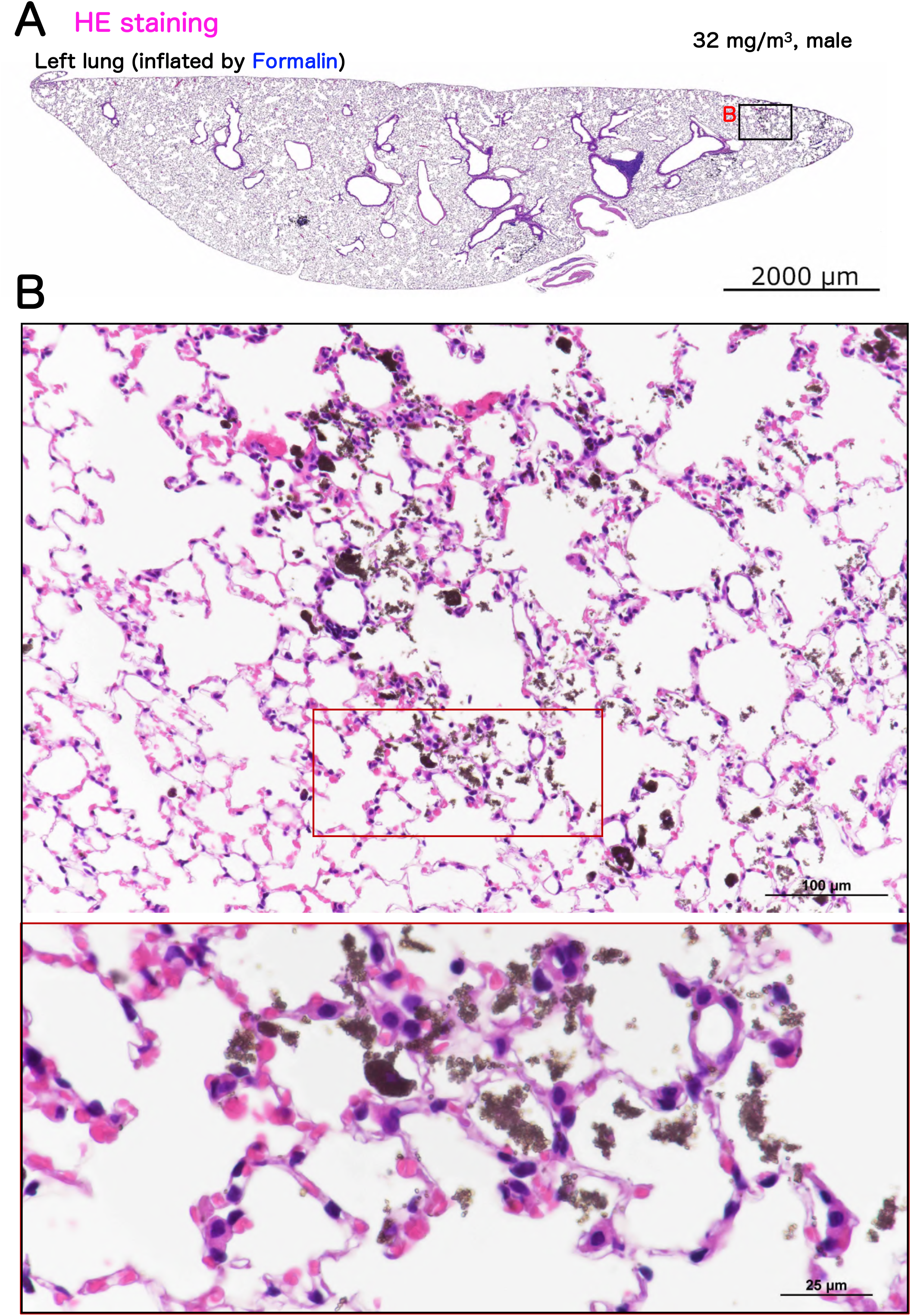
Representative microscopic photographs of male rasH2 mouse left lung after inhalation exposure to TiO_2_ NPs (32 mg/m^3^), same mouse of fig.2. For the left lung, formalin was injected into the lung through the bronchus and stained with HE. A typical loupe image (A) of the entire right lungs and magnified image of inflammatory focus (B) have been shown. The inflammatory foci observed in the right lung were scattered in the left lung due to the injection of formalin, making it difficult to observe them clearly.

**Figure S3.**
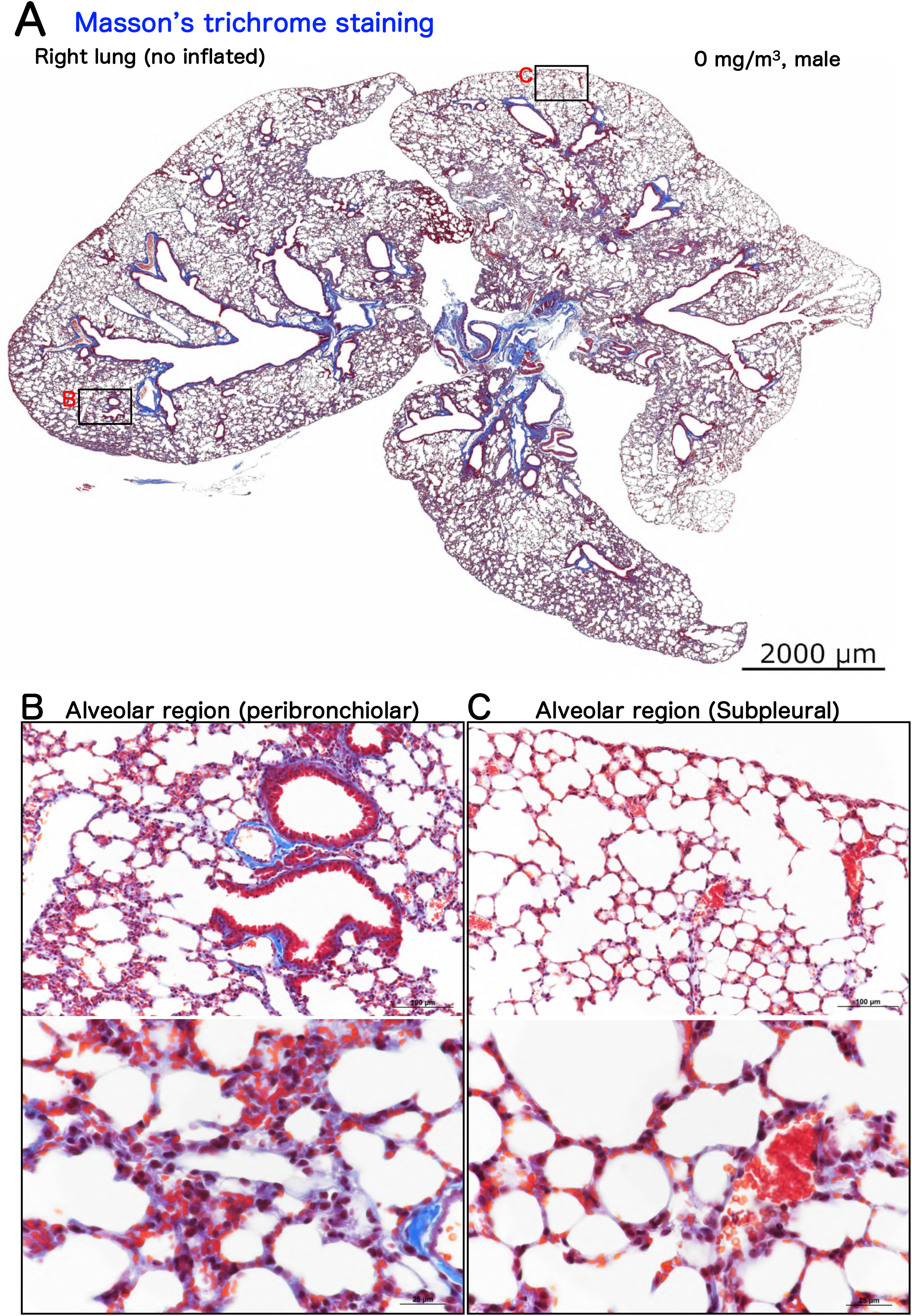
Representative microscopic photographs of Masson’s trichrome staining of male control rasH2 mouse right lungs. A typical loupe image (A) of the entire right lungs and magnified images of peribronchiolar (B) and subpleural (C) alveolar regions have been shown.

**Figure S4.**
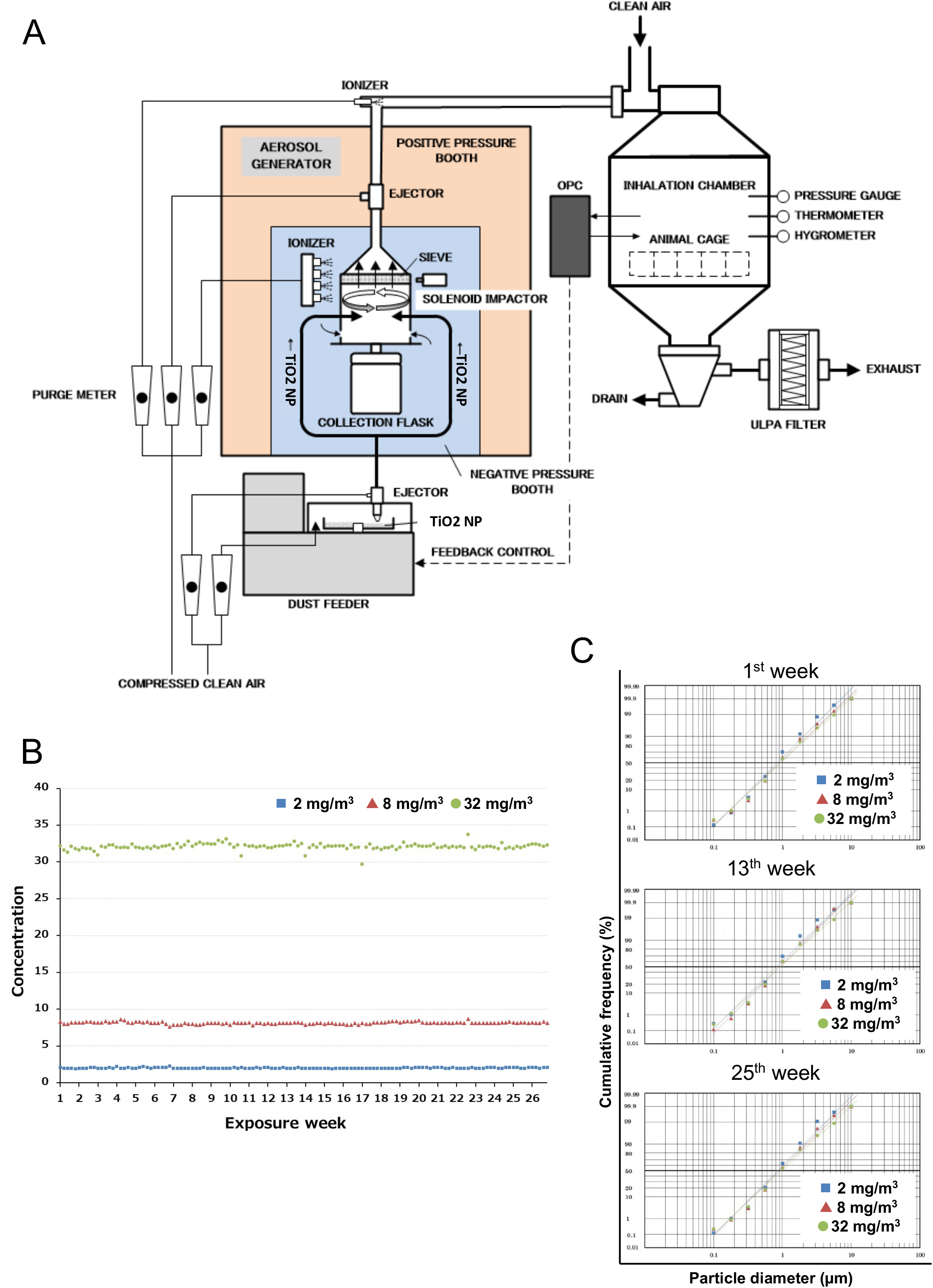
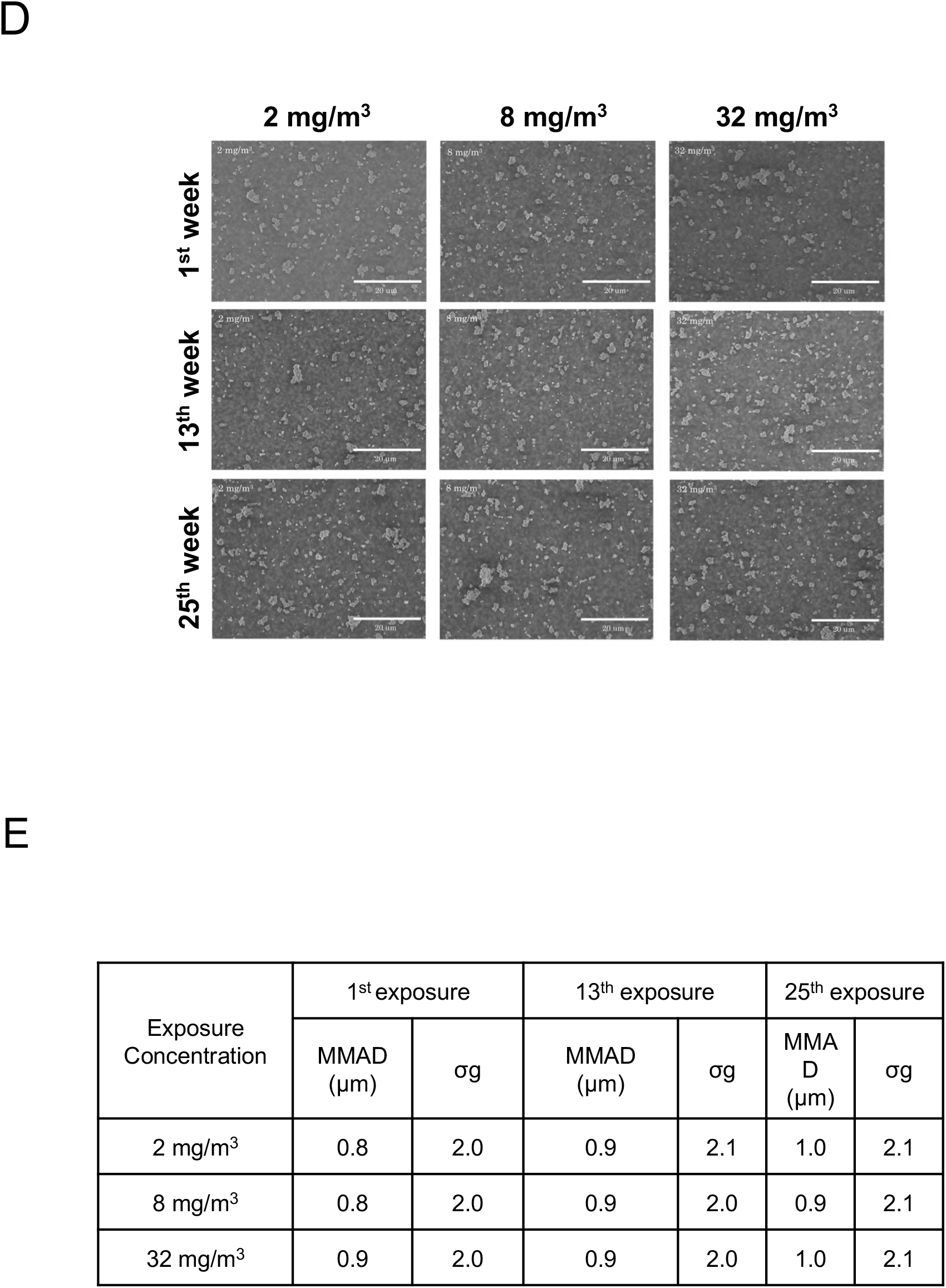
The whole body inhalation exposure system (A), the averaged TiO_2_ NPs concentration in the chamber per each exposure day (B), cumulative frequency distribution graphs with logarithmic probability (C), representative scanning electron microscope (SEM) images of the TiO_2_ NPs particles in the chambers (D) and Mass median aerodynamic diameter (MMAD) and geometric standard deviation (σg) in the chamber (E). Scale bar: 20 μm (panel D).

**Figure S5.**
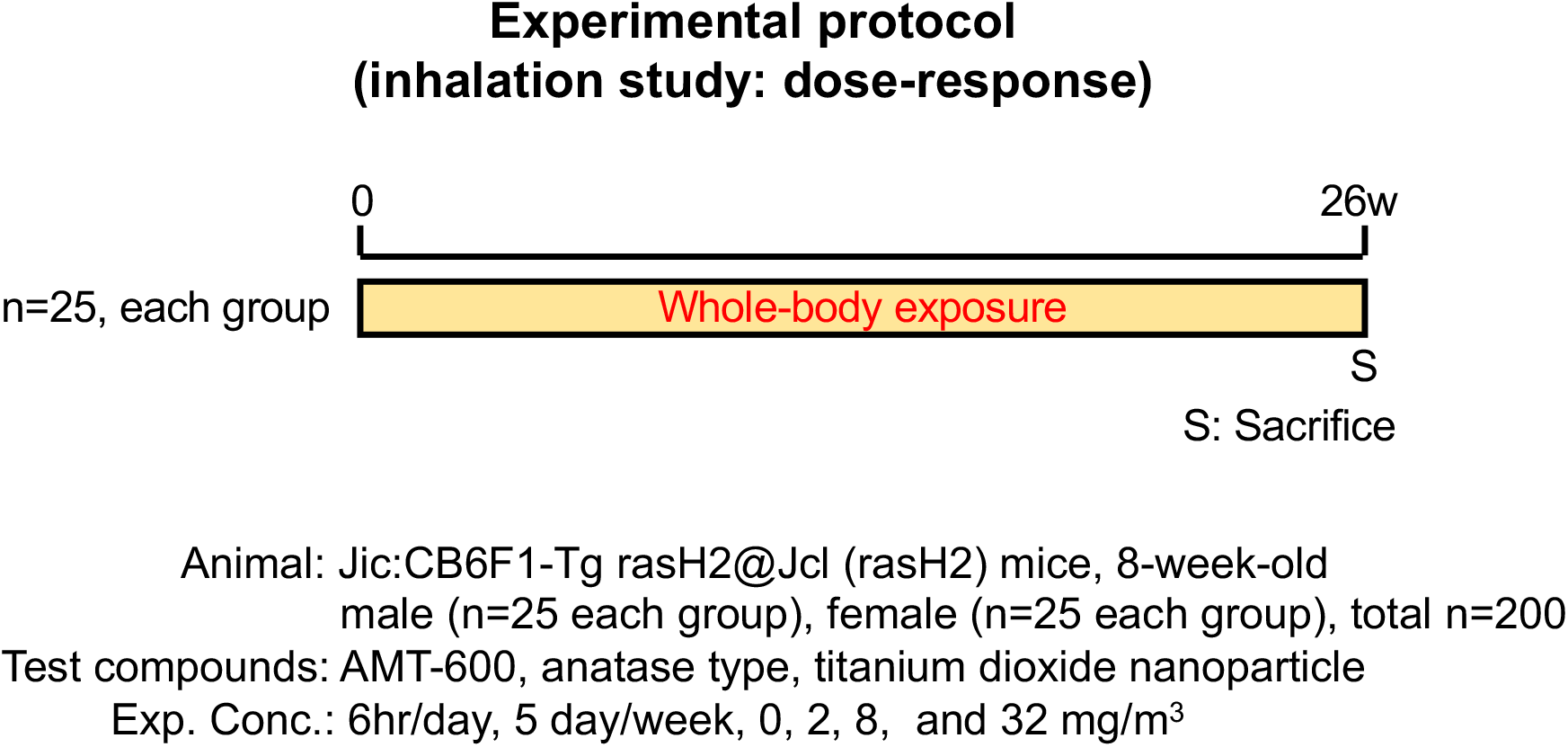
Design of animal experimental protocol used for this study.

**Table S1.** All blood-hematologic data observed in 26-week inhalation exposure study.

| Exposure concentration (mg/m3)<br>No. of Animals Exsaminated | Male |  |  |  | Female |  |  |  |
| --- | --- | --- | --- | --- | --- | --- | --- | --- |
|  | 0 | 2 | 8 | 32 | 0 | 2 | 8 | 32 |
|  | 24 | 23 | 25 | 25 | 24 | 25 | 23 | 23 |
| Hematology |  |  |  |  |  |  |  |  |
| RBCs (10<sup>6</sup>/µl) | 11.15±0.47 | 10.98±0.38 | 10.92±0.43* | 10.39±1.55** | 10.79±0.25 | 10.46±0.42** | 10.57±0.32 | 10.53±0.39* |
| Hemoglobin (g/dl) | 17.0±0.7 | 16.9±0.6 | 17.0±0.5 | 16.1±2.6 | 17.0±0.4 | 16.6±0.6 | 16.8±0.5 | 16.7±0.5 |
| Hmatocrit (%) | 51.6±1.5 | 50.9±1.6 | 51.0±2.1 | 49.0±7.0** | 50.7±1.0 | 49.9±1.2 | 50.1±1.5 | 50.2±1.8 |
| MCV (fl) | 46.3±1.3 | 46.3±0.7 | 46.7±1.0 | 47.2±1.5* | 47.0±0.6 | 47.7±1.2** | 47.4±0.7* | 47.7±0.6** |
| MCH (pg) | 15.3±0.2 | 15.3±0.3 | 15.6±0.4** | 15.4±0.6* | 15.8±0.3 | 15.9±0.3 | 15.9±0.3 | 15.9±0.3 |
| MCHC (g/dl) | 33.0±0.7 | 33.1±0.5 | 33.3±0.6 | 32.7±1.8 | 33.5±0.4 | 33.3±0.7 | 33.6±0.7 | 33.3±0.5 |
| Platelet (10<sup>3</sup>/µl) | 1334±121 | 1274± 69 | 1285± 93 | 1290±170 | 1166±100 | 1189±119 | 1122±163 | 1144± 61 |
| Reticulocyte (%) | 2.4±1.5 | 2.1±0.2 | 2.1±0.2 | 4.3±8.1 | 2.6±0.6 | 2.8±2.8 | 2.3±0.4 | 2.4±0.5 |
| WBC (10<sup>3</sup>/µl) | 1.86±0.84 | 1.59±0.84 | 1.67±1.18 | 1.58±0.81 | 2.58±2.10 | 2.35±2.08 | 1.98±1.16 | 1.72±1.71* |
| Differential WBC (%) |  |  |  |  |  |  |  |  |
| Neutrophil | 22± 7 | 24±11 | 22± 9 | 22± 7 | 26± 9 | 24±13 | 28±12 | 21± 9 |
| Lymphocyte | 63± 7 | 66±10 | 66±11 | 67± 9 | 67± 8 | 65±12 | 65±10 | 68±11 |
| Monocyte | 13±7 | 7±6** | 8±7 | 8±7* | 5±8 | 8±9 | 4±6 | 8±8 |
| Eosinophil | 2±1 | 2±1 | 3±1 | 2±1 | 2±1 | 2±1 | 3±1 | 2±1 |
| Basophil | 0±0 | 0±0 | 0±0 | 0±0 | 0±0 | 0±0 | 0±0 | 0±0 |
| Other | 0±1 | 0±0 | 0±0 | 0±0 | 0±0 | 0±0 | 0±0 | 0±0 |

**Table S2.** All blood-biochemistry data observed in 26-week inhalation exposure study.

| Exposure concentration (mg/m3)<br>No. of Animals Examined | Male |  |  |  | Female |  |  |  |
| --- | --- | --- | --- | --- | --- | --- | --- | --- |
|  | 0 | 2 | 8 | 32 | 0 | 2 | 8 | 32 |
|  | 23 | 23 | 25 | 25 | 24 | 25 | 23 | 23 |
| Biochemistry |  |  |  |  |  |  |  |  |
| Total protein (g/dl) | 5.1±0.1 | 5.3±0.2** | 5.3±0.2* | 5.2±0.2 | 5.2±0.2 | 5.4±0.2* | 5.4±0.2* | 5.4±0.2 |
| Albumin (g/dl) | 2.9±0.1 | 2.9±0.1 | 2.9±0.1 | 2.9±0.1 | 3.1±0.1 | 3.1±0.1 | 3.1±0.1 | 3.1±0.1 |
| A/G ratio | 1.3±0.1 | 1.2±0.1* | 1.2±0.1** | 1.2±0.1* | 1.4±0.1 | 1.3±0.1** | 1.3±0.1 | 1.4±0.1 |
| T-Bilirubin (mg/dl) | 0.06±0.01 | 0.06±0.01 | 0.05±0.01 | 0.05±0.01 | 0.05±0.01 | 0.04±0.01* | 0.05±0.01 | 0.06±0.02 |
| Glucose (mg/dl) | 201±26 | 231±20** | 236±29** | 223±20** | 190±25 | 202±25 | 219±26** | 216±27** |
| T-Cholesterol (mg/dl) | 74± 7 | 78± 8 | 76± 8 | 78±12 | 60± 7 | 64±11 | 64± 7 | 64± 7 |
| Triglyceride (mg/dl) | 45± 9 | 54±16 | 53±15 | 53±17 | 31±14 | 37±13 | 40±16 | 36±12 |
| Phospholipid (mg/dl) | 161±15 | 168±16 | 164±19 | 167±24 | 123±17 | 131±22 | 132±16 | 133±14 |
| AST (U/l) | 69±21 | 73±17 | 86±43 | 79±26 | 63±12 | 83±26** | 76±18 | 88±30** |
| ALT (U/l) | 26±15 | 26± 8 | 40±33 | 29±12 | 20±3 | 27±8** | 26±7** | 28±9** |
| LDH (U/l) | 250± 98 | 264± 65 | 293±122 | 260± 53 | 180±30 | 210±30** | 220±38** | 238±52** |
| ALP (U/l) | 205±18 | 211±20 | 205±12 | 209±21 | 332±34 | 323±39 | 321±44 | 338±41 |
| G-GTP (U/l) | 0.3±0.4 | 0.5±0.4 | 0.6±0.7 | 0.4±0.4 | 0.4±0.3 | 0.8±1.1 | 0.7±0.8 | 0.6±0.5 |
| CK (U/l) | 168±297 | 160±191 | 168±190 | 139±120 | 74±21 | 131±78** | 112±53** | 121±57** |
| Ureanitrogen (mg/dl) | 24.6±5.9 | 22.7±3.9 | 23.2±4.4 | 21.2±3.3 | 18.8±2.9 | 17.4±2.8 | 16.7±2.4* | 16.3±1.6** |
| Sodium (mEq/l) | 151±2 | 151±2 | 151±2 | 151±2 | 150±2 | 151±2 | 151±2 | 152±2** |
| Potassium (mEq/l) | 3.7±0.3 | 3.4±0.3** | 3.4±0.4* | 3.3±0.4** | 3.4±0.3 | 3.0±0.4** | 3.1±0.3* | 3.0±0.5** |
| Chloride (mEq/l) | 117±2 | 115±2** | 115±3 | 115±3 | 116±2 | 114±4 | 115±2 | 115±4 |
| Calcium (mg/dl) | 8.4±0.1 | 8.7±0.2** | 8.7±0.3** | 8.7±0.2** | 8.9±0.3 | 9.0±0.3 | 9.0±0.2 | 9.0±0.2 |
| Inorganic phosphrus (mg/dl) | 4.7±0.7 | 5.4±1.0* | 5.5±1.3* | 5.3±0.7* | 4.9±0.8 | 5.2±0.8 | 5.4±1.1 | 5.6±0.9 |

**Table S3.** Absolute organ weights observed in 26-week inhalation exposure study.

| Exposure concentration (mg/m3)<br>No. of Animals Exsaminated | Male |  |  |  | Female |  |  |  |
| --- | --- | --- | --- | --- | --- | --- | --- | --- |
|  | 0 | 2 | 8 | 32 | 0 | 2 | 8 | 32 |
|  | 25 | 24 | 25 | 25 | 24 | 25 | 24 | 23 |
| Organ (g) |  |  |  |  |  |  |  |  |
| Body Weight | 26.3±1.8 | 27.3±1.9 | 27.8±1.8* | 27.6±2.0 | 21.4±1.1 | 22.0±1.4 | 22.3±1.4 | 22.0±1.4 |
| Thymus | 0.056±0.019 | 0.054±0.012 | 0.054±0.012 | 0.056±0.013 | 0.049±0.013 | 0.054±0.022 | 0.054±0.011 | 0.054±0.014 |
| Adrenals | 0.012±0.002 | 0.012±0.003 | 0.012±0.003 | 0.014±0.004 | 0.014±0.003 | 0.015±0.003 | 0.015±0.002 | 0.015±0.004 |
| Testes/Ovaries | 0.261±0.029 | 0.253±0.039 | 0.264±0.020 | 0.266±0.024 | 0.030±0.006 | 0.040±0.013** | 0.037±0.007** | 0.038±0.007** |
| Heart | 0.175±0.016 | 0.175±0.014 | 0.181±0.024 | 0.177±0.017 | 0.145±0.011 | 0.149±0.012 | 0.143±0.010 | 0.141±0.011 |
| Lungs | 0.160±0.012 | 0.178±0.018** | 0.180±0.023** | 0.191±0.020** | 0.160±0.010 | 0.176±0.021** | 0.173±0.016** | 0.177±0.020** |
| Kidneys | 0.544±0.045 | 0.565±0.043 | 0.573±0.053 | 0.577±0.042 | 0.410±0.027 | 0.431±0.128 | 0.418±0.029 | 0.407±0.028 |
| Spleen | 0.066±0.015 | 0.065±0.010 | 0.085±0.086 | 0.102±0.096 | 0.080±0.012 | 0.087±0.016 | 0.087±0.016 | 0.081±0.013 |
| Liver | 1.170±0.090 | 1.244±0.110 | 1.297±0.121** | 1.282±0.121** | 1.009±0.078 | 1.071±0.104 | 1.103±0.093** | 1.071±0.090 |
| Brain | 0.472±0.013 | 0.479±0.011 | 0.476±0.016 | 0.483±0.016 | 0.496±0.015 | 0.502±0.014 | 0.504±0.022 | 0.498±0.018 |

**Table S4.** Relative organ weights observed in 26-week inhalation exposure study.

| Exposure concentration (mg/m3)<br>No. of Animals Exsaminated | Male |  |  |  | Female |  |  |  |
| --- | --- | --- | --- | --- | --- | --- | --- | --- |
|  | 0 | 2 | 8 | 32 | 0 | 2 | 8 | 32 |
|  | 25 | 24 | 25 | 25 | 24 | 25 | 24 | 23 |
| Organ (%) |  |  |  |  |  |  |  |  |
| Thymus | 0.211±0.069 | 0.196±0.040 | 0.193±0.043 | 0.201±0.036 | 0.230±0.061 | 0.248±0.103 | 0.240±0.046 | 0.245±0.068 |
| Adrenals | 0.045±0.009 | 0.043±0.011 | 0.044±0.011 | 0.049±0.014 | 0.064±0.014 | 0.068±0.013 | 0.068±0.010 | 0.070±0.016 |
| Testes/Ovaries | 0.995±0.130 | 0.933±0.161 | 0.953±0.093 | 0.971±0.099 | 0.141±0.025 | 0.179±0.054** | 0.165±0.030* | 0.173±0.030** |
| Heart | 0.664±0.053 | 0.645±0.051 | 0.649±0.076 | 0.645±0.058 | 0.679±0.037 | 0.677±0.054 | 0.641±0.044** | 0.641±0.038** |
| Lungs | 0.608±0.042 | 0.654±0.055* | 0.645±0.066 | 0.696±0.071** | 0.749±0.041 | 0.799±0.088* | 0.775±0.054 | 0.808±0.079** |
| Kidneys | 2.068±0.138 | 2.074±0.114 | 2.063±0.165 | 2.102±0.172 | 1.920±0.096 | 1.953±0.528 | 1.871±0.110 | 1.855±0.106 |
| Spleen | 0.250±0.057 | 0.238±0.039 | 0.304±0.295 | 0.375±0.370 | 0.375±0.052 | 0.394±0.073 | 0.387±0.060 | 0.368±0.052 |
| Liver | 4.442±0.154 | 4.566±0.277 | 4.661±0.285** | 4.655±0.324** | 4.722±0.206 | 4.866±0.353* | 4.935±0.243* | 4.870±0.202 |
| Brain | 1.799±0.106 | 1.765±0.115 | 1.715±0.105 | 1.760±0.121 | 2.325±0.098 | 2.285±0.130 | 2.262±0.129 | 2.272±0.121 |

**Table S5.** Histopathological findings excluding lung and mediastinal lymph node observed in 26-week inhalation exposure study.

|  |  | Male |  |  |  | Female |  |  |  |
| --- | --- | --- | --- | --- | --- | --- | --- | --- | --- |
| Exposure concentration (mg/m<sup>3</sup>) |  | 0 | 2 | 8 | 32 | 0 | 2 | 8 | 32 |
| No. of Animals Examined |  | 25 | 25 | 25 | 25 | 25 | 25 | 25 | 25 |
| Histopathological findings |  |  |  |  |  |  |  |  |  |
| Nasal cavity |  |  |  |  |  |  |  |  |  |
|  | Exudate | 0 | 0 | 1 | 3 | 0 | 0 | 1 | 2 |
|  |  |  |  | <1> | <1> |  |  | <1> | <1> |
|  | Respiratory metaplasia, nasal gland | 25 | 15*** | 21 | 24 | 23 | 25 | 23 | 24 |
|  |  | <1> | <1> | <1> | <1> | <1> | <1> | <1> | <1> |
|  | Respiratory metaplasia, gland | 0 | 0 | 0 | 1 | 0 | 0 | 0 | 0 |
|  |  |  |  |  | <3> |  |  |  |  |
|  | Squamous metaplasia, respiratory epithelium | 0 | 0 | 0 | 0 | 0 | 0 | 0 | 1 |
|  |  |  |  |  |  |  |  |  | <1> |
|  | Eosinophilic change, olfactory epithelium | 13 | 9 | 18 | 15 | 16 | 14 | 16 | 19 |
|  |  | <1> | <1> | <1> | <1> | <1> | <1> | <1> | <1> |
|  | Eosinophilic change, respiratory epithelium | 19 | 13 | 23 | 23 | 0 | 0 | 0 | 0 |
|  |  | <1> | <1> | <1> | <1> |  |  |  |  |
| Liver |  |  |  |  |  |  |  |  |  |
|  | Angiectasis | 0 | 0 | 0 | 1 | 0 | 1 | 0 | 0 |
|  |  |  |  |  | <1> |  | <2> |  |  |
|  | Necrosis, focal | 1 | 0 | 2 | 2 | 0 | 0 | 1 | 1 |
|  |  | <1> |  | <1> | <1> |  |  | <1> | <1> |
|  | Fatty change, central | 0 | 0 | 1 | 0 | 0 | 0 | 0 | 0 |
|  |  |  |  | <1> |  |  |  |  |  |
|  | Focus of cellular alteration | 0 | 0 | 0 | 1 | 0 | 1 | 0 | 0 |
|  |  |  |  |  | <2> |  | <1> |  |  |
| Stomach |  |  |  |  |  |  |  |  |  |
|  | Hyperplasia, forestomach | 0 | 1 | 1 | 0 | 0 | 0 | 0 | 0 |
|  |  |  | <1> | <2> |  |  |  |  |  |
|  | Hyperplasia, glandular stomach | 0 | 0 | 1 | 0 | 0 | 0 | 0 | 0 |
|  |  |  |  | <2> |  |  |  |  |  |
|  | Erosion, forestomach | 0 | 0 | 0 | 0 | 0 | 1 | 0 | 0 |
|  |  |  |  |  |  |  | <1> |  |  |
| Pancreas |  |  |  |  |  |  |  |  |  |
|  | Hemorrhage | 0 | 0 | 1 | 0 | 0 | 0 | 0 | 0 |
|  |  |  |  | <1> |  |  |  |  |  |
| Kidney |  |  |  |  |  |  |  |  |  |
|  | Regeneration, proximal tubule | 1 | 0 | 0 | 0 | 0 | 0 | 0 | 0 |
|  |  | <1> |  |  |  |  |  |  |  |
|  | Hydronephrosis | 0 | 0 | 0 | 0 | 0 | 1 | 1 | 0 |
|  |  |  |  |  |  |  | <3> | <1> |  |
| Urinary bladder |  |  |  |  |  |  |  |  |  |
|  | Simple hyperplasia, urothelium | 1 | 0 | 0 | 0 | 0 | 0 | 0 | 0 |
|  |  | <1> |  |  |  |  |  |  |  |
| Spleen |  |  |  |  |  |  |  |  |  |
|  | Deposit of melanin | 1 | 0 | 2 | 3 | 6 | 3 | 6 | 1 |
|  |  | <1> |  | <1> | <1> | <1> | <1> | <1> | <1> |
|  | Extramedullary hematopoiesis | 1 | 0 | 0 | 2 | 1 | 1 | 1 | 0 |
|  |  | <1> |  |  | <1> | <1> | <1> | <1> |  |
| Pituitary gland |  |  |  |  |  |  |  |  |  |
|  | Cystic degeneration, anterior lobe | 0 | 1 | 0 | 0 | 0 | 0 | 0 | 0 |
|  |  |  | <1> |  |  |  |  |  |  |
|  | Rathke pouch | 0 | 0 | 0 | 0 | 1 | 0 | 0 | 0 |
|  |  |  |  |  |  | <1> |  |  |  |
| Thyroid |  |  |  |  |  |  |  |  |  |
|  | Ultimobranchial cyst | 0 | 0 | 0 | 0 | 0 | 0 | 1 | 0 |
|  |  |  |  |  |  |  |  | <1> |  |
| Parathyroid |  |  |  |  |  |  |  |  |  |
|  | Cyst | 2 | 0 | 3 | 0 | 0 | 1 | 0 | 1 |
|  |  | <1> |  | <1> |  |  | <1> |  | <1> |
| Adrenal |  |  |  |  |  |  |  |  |  |
|  | Spindle-cell hyperplasia | 21 | 18 | 20 | 22 | 25 | 25 | 25 | 25 |
|  |  | <1> | <1> | <1> | <1> | <1> | <1> | <1> | <1> |
| Testis |  |  |  |  |  |  |  |  |  |
|  | Tubular atrophy | 1 | 0 | 1 | 0 |  |  |  |  |
|  |  | <2> |  | <1> |  |  |  |  |  |
| Epididymis |  |  |  |  |  |  |  |  |  |
|  | Inflammatory cell infiltration | 0 | 0 | 1 | 0 |  |  |  |  |
|  |  |  |  | <1> |  |  |  |  |  |
|  | Debris of spermatic elements | 1 | 0 | 0 | 0 |  |  |  |  |
|  |  | <1> |  |  |  |  |  |  |  |
| Ovary |  |  |  |  |  |  |  |  |  |
|  | Thrombus |  |  |  |  | 0 | 1 | 0 | 0 |
|  |  |  |  |  |  |  | <2> |  |  |
|  | Cyst |  |  |  |  | 0 | 1 | 0 | 0 |
|  |  |  |  |  |  |  | <1> |  |  |
| Uterus |  |  |  |  |  |  |  |  |  |
|  | Cystic endometrial hyperplasia |  |  |  |  | 25 | 24 | 25 | 25 |
|  |  |  |  |  |  | <1> | <1> | <1> | <1> |
|  | Vascular anomaly |  |  |  |  | 1 | 0 | 0 | 0 |
|  |  |  |  |  |  | <1> |  |  |  |
| Peripheral nerve |  |  |  |  |  |  |  |  |  |
|  | Cyst | 0 | 0 | 0 | 0 | 0 | 0 | 0 | 1 |
|  |  |  |  |  |  |  |  |  | <1> |
| Cartilage |  |  |  |  |  |  |  |  |  |
|  | Necrosis, focal | 1 | 0 | 0 | 0 | 0 | 0 | 1 | 0 |
|  |  | <1> |  |  |  |  |  | <1> |  |
| Peritoneum |  |  |  |  |  |  |  |  |  |
|  | Cyst | 0 | 0 | 0 | 0 | 0 | 0 | 1 | 0 |
|  |  |  |  |  |  |  |  | <1> |  |

Note: Values indicate number of animals bearing lesions. The value in angle bracket indicate the average of severity grade index of the lesion. The average of severity grade is calculated with a following equation. $\Sigma(\text{grade} \times \text{number of animals with grade}) / \text{number of affected animals}$ . Grade: 1, slight; 2, moderate; 3, marked; 4, severe. Significant difference: \*, p<0.05; \*\*, p<0.01; \*\*\*, p<0.001 by Chi square test compared with each cont.

